# Efficient multi-fidelity computation of blood coagulation under flow

**DOI:** 10.1101/2023.05.29.542763

**Authors:** Manuel Guerrero-Hurtado, Manuel Garcia-Villalba, Alejandro Gonzalo, Pablo Martinez-Legazpi, Andy M. Kahn, Elliot McVeigh, J. Bermejo, Juan C. del Alamo, Oscar Flores

## Abstract

Clot formation is a crucial process that prevents bleeding, but can lead to severe disorders when imbalanced. This process is regulated by the coagulation cascade, a biochemical network that controls the enzyme thrombin, which converts soluble fibrinogen into the fibrin fibers that constitute clots. Coagulation cascade models are typically complex and involve dozens of partial differential equations (PDEs) representing various chemical species’ transport, reaction kinetics, and diffusion. Solving these PDE systems computationally is challenging, due to their large size and multi-scale nature.

We propose a multi-fidelity strategy to increase the efficiency of coagulation cascade simulations. Leveraging the slower dynamics of molecular diffusion, we transform the governing PDEs into ordinary differential equations (ODEs) representing the evolution of species concentrations versus blood residence time. We then Taylor-expand the ODE solution around the zero-diffusivity limit to obtain spatiotemporal maps of species concentrations in terms of the statistical moments of residence time, 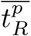, and provide the governing PDEs for 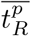. This strategy replaces a high-fidelity system of *N* PDEs representing the coagulation cascade of *N* chemical species by *N* ODEs and *p* PDEs governing the residence time statistical moments. The multi-fidelity order(*p*) allows balancing accuracy and computational cost, providing a speedup of over *N/p* compared to high-fidelity models.

Using a simplified coagulation network and an idealized aneurysm geometry with a pulsatile flow as a benchmark, we demonstrate favorable accuracy for low-order models of *p* = 1 and *p* = 2. These models depart from the high-fidelity solution by under 16% (*p* = 1) and 5% (*p* = 2) after 20 cardiac cycles.

The favorable accuracy and low computational cost of multi-fidelity models could enable unprecedented coagulation analyses in complex flow scenarios and extensive reaction networks. Furthermore, it can be generalized to advance our understanding of other systems biology networks affected by blood flow.

## Introduction

Blood coagulation, or clotting, is a highly regulated mechanism vital in sealing wounded blood vessels to prevent bleeding. Abnormal or excessive clotting can result in serious medical conditions, such as stroke and deep vein thrombosis. Therefore, blood coagulation has been the subject of extensive research, and understanding its mechanisms is crucial to diagnose and manage numerous diseases.

The initiation of blood coagulation is an enzymatic cascade that amplifies thrombin concentration in plasma, activating fibrin polymerization to form a clot [10, 39]. This cascade can be started via extrinsic and intrinsic pathways. The extrinsic pathway is triggered by vessel injury-mediated release of coagulation factor VII (FVII) and tissue factor (TF) into the bloodstream. The intrinsic pathway is auto-initiated, i.e., it does not require exposure to an extravascular tissue factor, and begins with the activation of plasma factors XII, XI, IX, and VIII. Both pathways eventually activate factor X, converging into the common pathway that amplifies thrombin, which in turn converts fibrinogen into fibrin filaments and activates factor XIII, which cross-links the fibrin mesh [22, 50]. The initiation, propagation, and inhibition of this complex process involve a network of over 80 biochemical reactions [52].

Since thrombosis is a ubiquitous complication of cardiovascular diseases and device implantation [46, 59], there is an abundance of computational models considering coagulation in diverse physiological and anatomical settings. Intra-luminal thrombogenesis in an abdominal aortic aneurysm has been studied using an 18-equation model based on tissue factor activation [28, 34], providing an integrated mechanochemical picture of the process [6]. The same model was used later to study thrombogenesis in an infarcted left ventricle [53]. More complex models have also been used in the literature, either increasing the number or reaction equations involved in the model [25], or including the effects of issue factor, platelet activation, and clot porosity on thrombus growth [35]. Other models have analyzed the contribution of intrinsic and extrinsic pathways at different timescales, highlighting the multi-stage character of the coagulation process [44].

Eulerian-Lagrangian approximations can be used to integrate coagulation cascade reaction-advection-diffusion equations with platelet activation and deposition. Researchers have used this approach to describe the evolution of non-activated and activated platelet concentrations [21], using the simulated velocity fields to track platelet activation and accumulation. The equations were solved using a stabilized finite element method. The accumulation model [33] accounts for various factors, including plasma-phase and membrane-phase reactions, coagulation inhibitors, and the presence of activated and unactivated platelets. Other groups have used similar strategies, like coupling a calibrated platelet aggregation model, which accounts for adhesion forces between platelet-platelet and platelet-wall at low and high shear rate levels, with an extrinsic coagulation cascade initiation model [61]. In this case, the coagulation cascade was based on a model using 23 chemical species [2].

The large number of coupled partial differential equations (PDEs) representing the reaction, advection, and diffusion of the species involved in the coagulation cascade creates stringent requirements for numerical simulation. This problem is aggravated by the high numerical cost of solving each PDE, owing to the disparate timescales associated with the flow, reaction kinetics, and diffusion [14, 38, 60]. The dimensional parameters involved in these processes for mid-size arteries are: flow velocity, *U*_*c*_ ∼ 10cm/s; vessel diameter, *L*_*c*_ ∼ 1cm; the cardiac cycle’s period, *t*_*c*_ ≈ 1*s*; timescales of enzymatic reaction kinetics, ranging from a few seconds to hundreds of seconds (*t*_*r*_ ∼ 10^2^s) [4, 28, 38] and mass diffusivity coefficients for species in blood, *D*_*i*_ ∼ 10^−6^cm^2^/s [6, 47, 62]. The relatively slow reaction times in these systems require running simulations over many cardiac cycles to reach convergence, i.e., *t*_*r*_ ≫ *t*_*c*_. Moreover, the slow diffusion of reactive species creates extremely thin boundaries in their concentration fields since the Peclet number, *Pe* = *U*_*c*_*L*_*c*_*/D*_*i*_ = *t*_*d*_*/t*_*a*_ ≈ 10^7^, which measures the ratio between convective and diffusive transport, is very large. For reference, the Schmidt number, which measures the ratio between the viscous diffusion acting in the Navier-Stokes equations and the mass diffusivity acting in the coagulation system’s equations, is *Sc* = *ν/D*_*i*_ ∼ 10^4^ where *ν* ≈ 4 × 10^−2^cm^2^/s is the kinematic viscosity of blood. Consequently, the spatial discretization of the coagulation system’s equations requires much finer computational grids than the ones used to solve the Navier-Stokes equations. This problem is well described in the computational fluid dynamics (CFD) literature, and it is common to many non-reactive cardiovascular transport problems [29]. Previous simulations of the coagulation cascade under flow have proposed concessions to reduce computational cost. First, the multi-scale nature of reaction kinetics have been simplified by assuming that fast-reacting species are in equilibrium, leading to reduced models with fewer chemical species [62], or by using phenomenological models [5]. The latter have been used to investigate hypercoagulability in the left heart by considering fibrin production from fibrinogen and thrombin [45]. Second, in most if not all studies, the Peclet number has been decreased explicitly by prescribing unphysically high values for *D*_*i*_ [6, 53], or implicitly by using a diffusive numerical discretization (e.g., upwinding first-order finite differences). In non-reactive transport problems, this concession has been justified on the basis of accounting for additional noise sources and as long as the effective *Pe* remains ≫ 1 [17]. However, its adequacy is more questionable in reacting problems like the coagulation cascade, where the reaction timescale is intermediate between the transport and diffusive timescales. Another compromise adopted by some authors is to shorten the time integration to a few cardiac cycles, focusing only on the initial phases of thrombin activation [53]. Finally, many studies have considered idealized two-dimensional geometries to save computational cost [6, 14, 35, 38, 45]. In summary, there is an unmet need for computationally efficient strategies to model the coagulation cascade under flow.

We introduce a multi-fidelity modeling approach to significantly reduce the computational cost of coagulation cascade simulations in flowing blood. This approach transforms the reaction-advection-diffusion equations for species concentrations (*u*_*i*_, *i* = 1, …, *N*) into a system of ODEs by using blood residence time 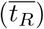 as the independent variable. The resulting model requires integrating only one PDE for 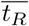 and *N* ODEs for the coagulatory species, whose concentration fields can be mapped as 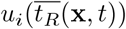. The transformation is exact for zero diffusivity. For small, finite diffusivity, the model can be Taylor-expanded in terms of the residence time statistical moments, i.e., 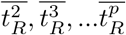, to derive a family of customizable, multi-fidelity models that offer a balance between cost and accuracy. We compare 1st-order (MuFi-1: 1 PDE, *N* ODEs) and 2nd-order (MuFi-2: 2 PDEs, *N* ODEs) multi-fidelity models with the high-fidelity (HiFi: *N* -PDEs) model for pulsatile flow through an aneurysm-like geometry, using a three-species coagulation system [62] as a benchmark. The MuFi-1 and HiFi models show good agreement up to *t* = 10*t*_*c*_ cardiac cycles, while this agreement is extended to *t* = 18*t*_*c*_ cycles for MuFi-2. The proposed family of multi-fidelity coagulation models could benefit researchers in the field by enabling them to simulate and analyze complex blood coagulation phenomena more quickly and accurately, thus advancing our understanding of the underlying mechanisms and informing clinical practice.

## Methods

This section presents a multi-fidelity model to reduce the cost of simulating coagulation networks of *N* species in flowing blood. To facilitate the model’s presentation, we first review the standard high-fidelity model (*N* -PDE) and introduce its first-order (1-PDE, *N* -ODE) approximation, which neglects diffusion. We then introduce the high-order approximation (2-PDE, *N* -ODE), define our benchmark flow problem and coagulation reaction system, and describe the numerical discretization methods.

### High-Fidelity and First-order Multi-Fidelity Coagulation Models

We consider blood as a continuum in space and time (**x**, *t*) flowing with velocity **v**(**x**, *t*), and model its coagulation by the system of reaction-advection-diffusion equations

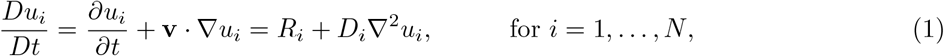

where *D/Dt* denotes material derivative, the subindex *i* indicates each of the *N* species involved in coagulation system, *u*_*i*_(**x**, *t*) its concentration field, *R*_*i*_(*u*_1_, *u*_2_,…, *u*_*N*_) its reaction rate from chemical kinetics, and *D*_*i*_ its diffusivity coefficient. This system of *N* PDEs is denoted the high-fidelity (HiFi) model. Assuming that **v**(**x**, *t*) is known, this HiFi model can be solved with some appropriate initial and boundary conditions for *u*_*i*_. For simplicity, we consider uniform initial conditions *u*_*i*_(**x**, 0) = *u*_*i*,0_, Dirichlet boundary conditions at the domain flow inlets (*u*_*i*_ = *u*_*i*,0_), and homogeneous Neumann boundary conditions (*∂u*_*i*_/*∂n* = 0) at solid surfaces and flow outlets.

The eqs. (1) can be written in non-dimensional form using the flow velocity scale *U*_*c*_ and vessel length scale *L*_*c*_,

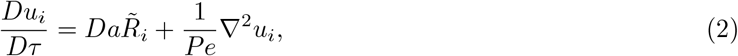

where *τ* = *tU*_*c*_*/L*_*c*_ is a dimensionless time variable, 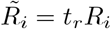is a dimensionless reaction rate, the Damköhler number *Da* = *L*_*c*_/(*t*_*r*_*U*_*c*_) measures the relative importance of reaction kinetics 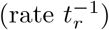 and convective terms, and the Péclet number *Pe* = *U*_*c*_*L*_*c*_*/D*_*i*_ measures the relative importance of convection over diffusion. Using typical values corresponding to mid-size arteries and the reaction rate and diffusivity of coagulation cascade species (i.e. *U*_*c*_ ∼ 10 cm/s, *L*_*c*_ ∼ 1 cm, *t*_*r*_ ∼ 10^2^ s, *D*_*i*_ ∼ 10^−6^ cm^2^/s) yields *Da* ∼ 10^−3^ and 1*/Pe* ∼ 10^−7^, so both terms on the right-hand side of eq. (2) are small. However, the reaction term is the only forcing in the equation and cannot be neglected, while the diffusive term is even smaller and can be neglected. Doing so simplifies eq. (1) to

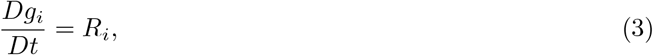

where *g*_*i*_ approximates *u*_*i*_ in the limit of zero molecular diffusivity. The simplified model has no mixing, making the reaction rate within each fluid element independent of the species concentrations in surrounding elements, and exclusively dependent on its *age*. Consequently, it should be possible to write an ODE system for the coagulation system of each fluid element in a Lagrangian frame that follows the element as it moves with the flow. To avoid the complication of tracking Lagrangian trajectories, we follow previous works [51] and resort to the PDE governing the residence time

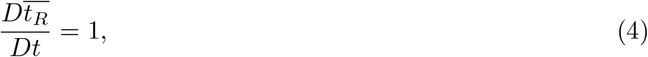

which can be used to calculate the *age* of fluid elements in the region of interest. Applying the chain rule on eq. (3) and taking into account eq. (4), we obtain

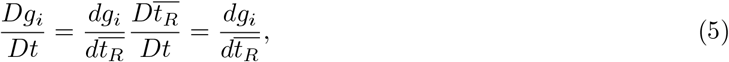

which simplifies the HiFi PDE system eqs. (1) into the ODE system

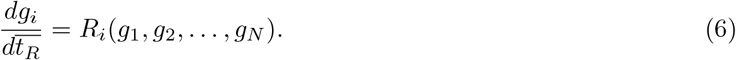

We note that this ODE system is the same for all fluid elements, as it has no explicit dependence on **x**, and each fluid element’s pathline information is implicitly encoded by the spatial dependence of 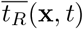. This model involves solving a system of *N* ODEs (i.e., eq. (6) for *i* = 1 …*N*) to obtain 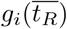, solving one PDE to calculate 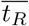, and mapping 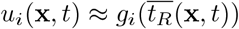. Because it combines solving ODEs and PDEs, we consider this approximate model a multi-fidelity (MuFi) model. When *N* is large, the MuFi model can be significantly cheaper to run than the HiFi model, which involves solving *N* PDEs.

### Higher-order Multi-Fidelity Approximations

In the previous section, we used overline notation for residence time to emphasize that 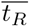 as defined in eq. (4) is the *ensemble* average age of all molecules within a fluid element, defined as the first integral moment of its probability density function, *f*_*T*_ :

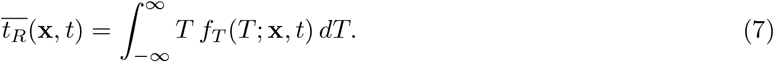

Considering residence time as a stochastic variable is important because, as discussed in the Introduction, the numerical solutions to transport PDEs like eq. (4) explicitly include unphysically large diffusivities or employ discretization methods that implicitly introduce numerical diffusivity. Numerical dissipation helps control instabilities and spurious oscillations but it is a source of numerical error [24]. This error makes 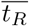 satisfy an equivalent differential equation (EDE) instead of the theoretical PDE given by eq. (4). The specific form of the EDE depends on the details of the temporal integration scheme and the approximation used to discretize the spatial derivatives. For commonly used, first-order, dissipative methods, the EDE for 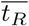 takes the form

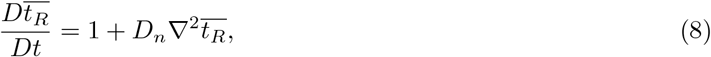

where the coefficient *D*_*n*_ ∼ *c*Δ*x*/2 represents the diffusivity of the numerical method, where *c* is the flow characteristic velocity and Δ*x* the mesh spatial resolution [24].

In this *effective* scenario, diffusivity creates uncertainty in the residence time. This phenomenon can be shown using Itô’s differentiation [26] to derive the EDE for the second-order moment of the residence time, 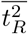, as

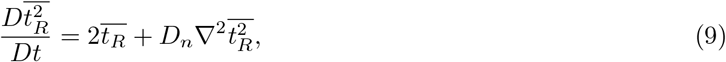

where

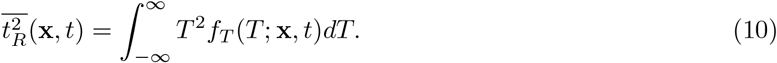

Then, it is possible to express this equation in terms of the residence time variance, 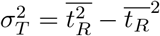, as

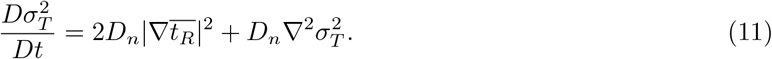

The interested reader can find the derivation of eqs. (8) and (11) in S1 Appendix. Of note, when *D*_*n*_ = 0 and *f*_*T*_ is a Dirac delta function, eqs. (8) and (9) 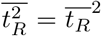 and zero residence time variance, i.e., 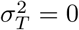. But when *D*_*n*_ ≠ 0, the non-negative forcing term in eq. (11) causes 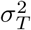 to increase unless residence time is constant. Note also that for higher order numerical methods, the differential operator multiplying *D*_*n*_ in the EDEs (8) and (9) will involve higher order derivatives, with a dispersive or dissipative character depending on the order of the temporal and spatial discretizations. This will result in more complex evolution equations for 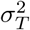, without changing the fact that numerical diffusion results in an increase on 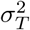 in regions with strong gradients of 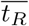.

The growth of 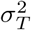causes errors in the MuFi model derived in the previous section as time increases. To illustrate these errors and derive higher-order corrections, it is convenient to express the concentration of chemical species as

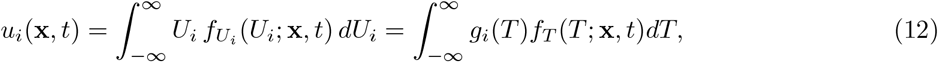

where *U*_*i*_ is the concentration’s statistical variable, 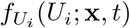 is its probability density function, and *f*_*T*_ (*T* ; **x**, *t*) is the probability density function of the residence time. The substitution *U*_*i*_ = *g*_*i*_(*T*) is warranted because, in the absence of diffusion, *σ*_*T*_ is zero and *f*_*T*_ is a Dirac delta, yielding 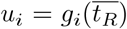, consistent with the definition of *g*_*i*_ in eq. (3).

Assuming next that *g*_*i*_(*T*) is second-order differentiable, we can Taylor-expand it around 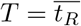 and integrate the expansion to obtain

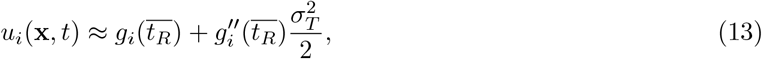

where the primes denote derivatives. This result demonstrates that the MuFi model 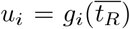error grows with the residence time variance for non-linear reaction systems (i.e., those with *g*^*ll*^ ≠ 0). But, more important, it provides a high-order correction that is also an inexpensive MuFi model as it only requires solving one additional PDE.

In summary, we introduce two multi-fidelity models that balance cost with accuracy:

- **First-order (MuFi-1)**:
- Solve N ODEs eqs. (6) to calculate *g*_*i*_(*t*).
- Solve one PDE eq. (4) to calculate 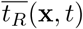
- Map 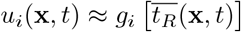.
- **Second-order (MuFi-2)**:
- Solve N ODEs eqs. (6) to calculate *g*_*i*_(*t*) and its second temporal derivative 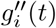.
- Solve two PDEs: Calculate 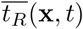 from eq. (4) and 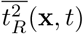 from

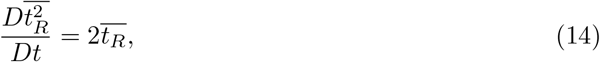

which follows from ignoring the numerical diffusion term from the EDE in eq. (9).

- Calculate 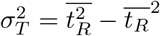, and map 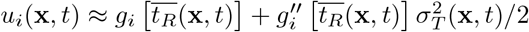.

This procedure can be extended to higher orders of approximations by retaining additional terms in the Taylor expansion of *g*_*i*_ around 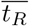, and solving additional PDEs for higher-order moments of the residence time. For example, the MuFi model of third order (involving N-ODEs and three PDEs) is derived in Appendix. Finally, we note that the PDEs to be solved in the MuFi-2 model are the *true* PDEs (i.e., eqs. (4) and (14)), not the EDEs (eqs. (8) and (9)). If the PDEs explicitly include diffusivity, then a diffusive term should be added. If not, the resulting 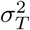will capture the effect of the discretization’s numerical diffusivity, *D*_*n*_, regardless of the order of the numerical method employed to solve eqs. (4) and (14). We emphasize that explicit knowledge of *D*_*n*_ is not required, which is advantageous since this diffusivity may be constant or vary in space and time depending on the numerical discretization.

### Computational cost estimates

We estimate the computational cost of running a HiFi model in *D* dimensions to be *αNn*^*D*+1^ floating point operations (FLOPs), where *N* is the number of species, *α* ∼ *O*(10^2^) is a parameter that depends on the numerical discretization scheme and the function evaluations needed to calculate the reaction rates, and *n* is the number of elements in the spatial mesh along each direction. Note that the exponent *D* + 1 reflects that the temporal resolution is linked to the spatial resolution by the CFL condition to guarantee an accurate temporal integration. As a consequence the number of time steps required to reach a finite integration time is proportional to *n*. On the other hand, a MuFi-p model is estimated to require *βNn* + *θpn*^*D*+1^ FLOPs, where the first term is the cost of integrating the *N* ODEs for the species reaction kinetics and the second term is the cost of solving the *p* PDEs governing the statistical moments of residence time. Like in the HiFi model, the parameters *β* and *θ* depend on the numerical implementation and the right-hand-side terms in the governing equations. It is worth noting that *θ* < *α* because the forcing terms in the residence time equations (see e.g., eqs. 4 and14) are simpler than the reaction rate terms in the equations governing species concentration (see, e.g., eqs. 18). Based on these estimates, one can see that MuFi models achieve a speedup ∼ (*α/θ*) (*N/p*) > *N/p* as soon as *n* is moderately large. Furthermore, MuFi models significantly reduce the memory allocation necessary to run simulations, which could lead to additional speedups in parallel implementations by avoiding the overheads associated with message passing and loss of cache memory coherence.

### Test Case: Coagulation Cascade in an Idealized Aneurysm

To compare the multi-fidelity models MuFi-1 and MuFi-2 with the high-fidelity model based on the PDE system (1), we considered a simplified coagulation cascade model under pulsatile flow through an idealized two-dimensional geometry (Fig 1). This flow geometry broadly resembles a cerebral aneurysm or the left atrial appendage, two cardiovascular sites associated with thrombosis [1, 9, 19, 42, 57] The parent vessel is modeled as a straight tube of diameter *H*, and the aneurysm is modeled as a circular cavity of radius 0.75*H*. The center of the cavity is located such that the aneurysm neck size is *H*. The corners at the aneurysm neck are smoothed with a radius of curvature equal to 0.067*H*, to avoid sharp corners in the geometry. The pulsatile flow was driven by imposing a two-dimensional Womersley flow as the inflow boundary condition (see S3 Appendix). The Reynolds and Womersley numbers are *Re* = *U*_*c*_*H/ν* = 500 and 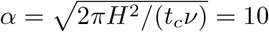, respectively, where *U*_*c*_ is the maximum velocity. These values of *Re* and *α* are representative of intracranial saccular aneurysms [32]. Fig 2 shows the time history of the mass flow rate and the inlet velocity profiles at two time instants, corresponding to the minimum (*t/t*_*c*_ = 0.5) and maximum (*t/t*_*c*_ = 1) flow rates through the vessel.

**Figure 1:**
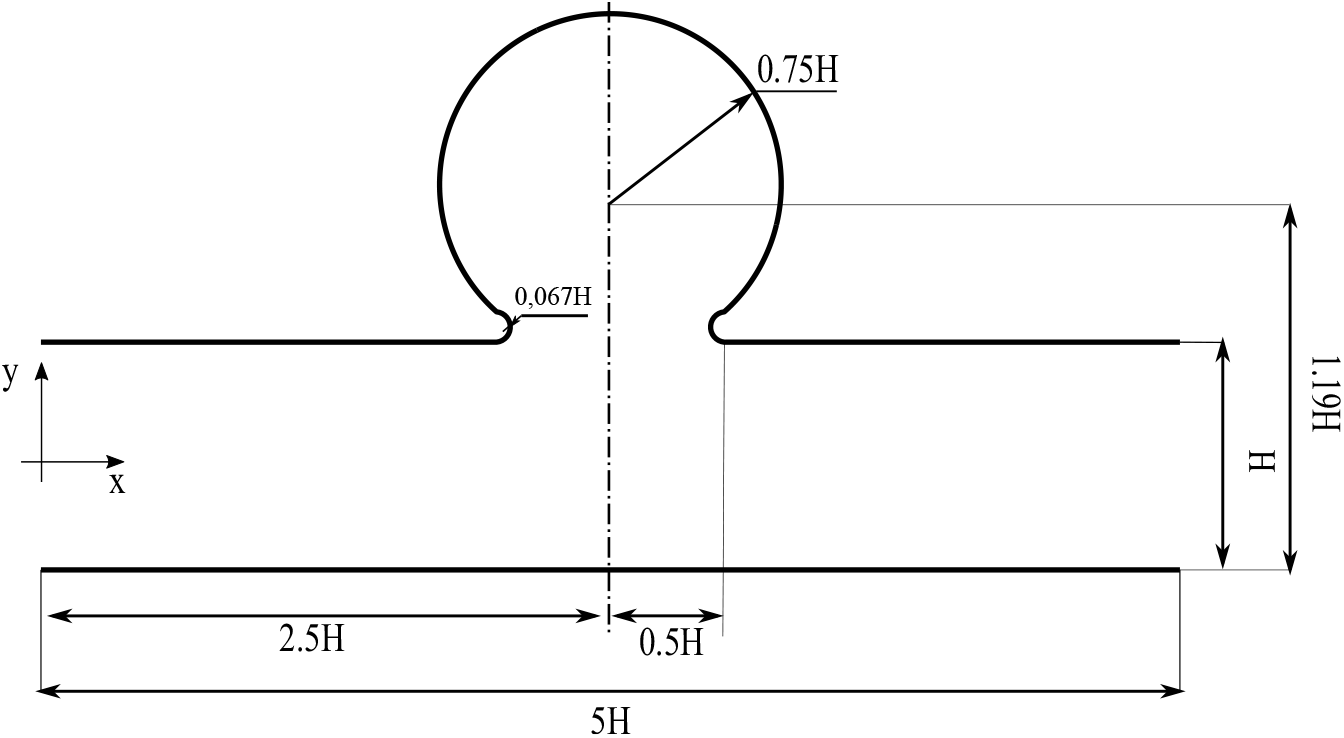
Idealized aneurysm geometry.

**Figure 2:**
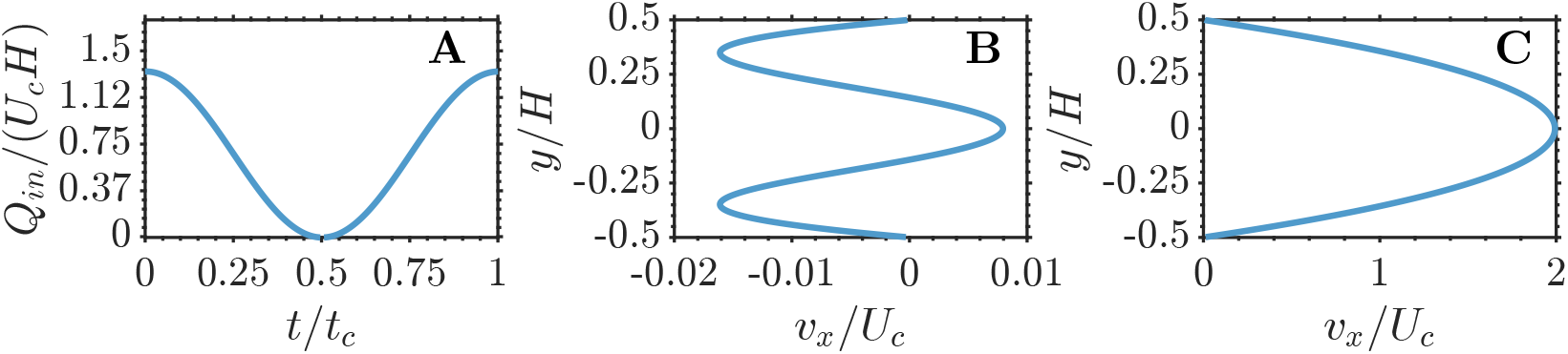
Womersley flow profiles. A: Time evolution of the mass flow rate through the vessel. B and C: Velocity profile at the inlet at *t* = 0.5*t*_*c*_ and *t* = *t*_*c*_.

To model the coagulation cascade, we chose a three-state system considering thrombin, factor XIa, and activated protein C (PCa) [62]. This simplified system was derived from a classic nine-state system (XIa, IXa, XA, tissue factor, prothrombin, thrombin, VIIIa, Va, and PCa) under the assumption that faster reactions are in instantaneous equilibrium. The non-dimensional concentrations of thrombin (*u*_1_), factor XIa (*u*_2_) and PCa (*u*_3_) are defined as

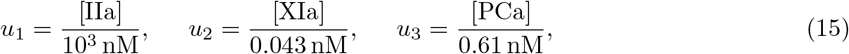

and the corresponding source terms in eq. (1) are

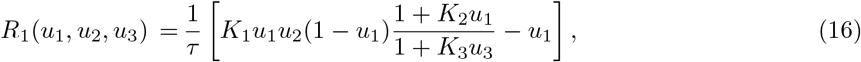

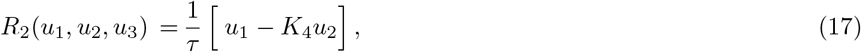

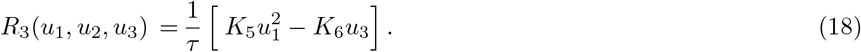

In these equations, thrombin production is activated by thrombin and factor XIa, and inhibited by PCa. In turn, thrombin activates the production of both factor XIa and PCa, thereby establishing a positive and a negative feedback loop. Finally, the non-dimensional thrombin concentration, *u*_1_, is bounded by one, whereas the concentrations of factor XIa (*u*_2_) and PCa (*u*_3_) are not [30, 62]. The values of the non-dimensional reaction rates of the model (*K*_*i*_) and their characteristic time-scale (*τ*) are taken from previous works [14], and reported in table 1.

**Table 1:** Coagulation model parameters.

| $K_1$ | $K_2$ | $K_3$ | $K_4$ | $K_5$ | $K_6$ | $\tau/t_c$ |
| --- | --- | --- | --- | --- | --- | --- |
| 6.85 | 20.0 | 2.36 | 0.087 | 15.0 | 0.04 | 25.8 |

Fig. 3 shows the evolution of *u*_*i*_ (*i* = 1, … 3) in stagnant blood (i.e. **v** = **0**) with uniform initial concentrations *u*_1,0_ = 0.0604, *u*_2,0_ = 0.0865 and *u*_3,0_ = 0.0975. These conditions produce a slow build up of thrombin and factor XIa over the initial 10 − 15 cardiac cycles, followed by rapid growth over the subsequent 5 cycles. At *t* ≈ 20*t*_*c*_ the nondimensional thrombin concentration flattens as it approaches unity. Although not shown in the figure, for longer times the growth of the inhibitor eventually brings down the concentrations of thrombin and factor XIa, driving the whole system to the equilibrium state *u*_*i*_ = 0.

**Figure 3:**
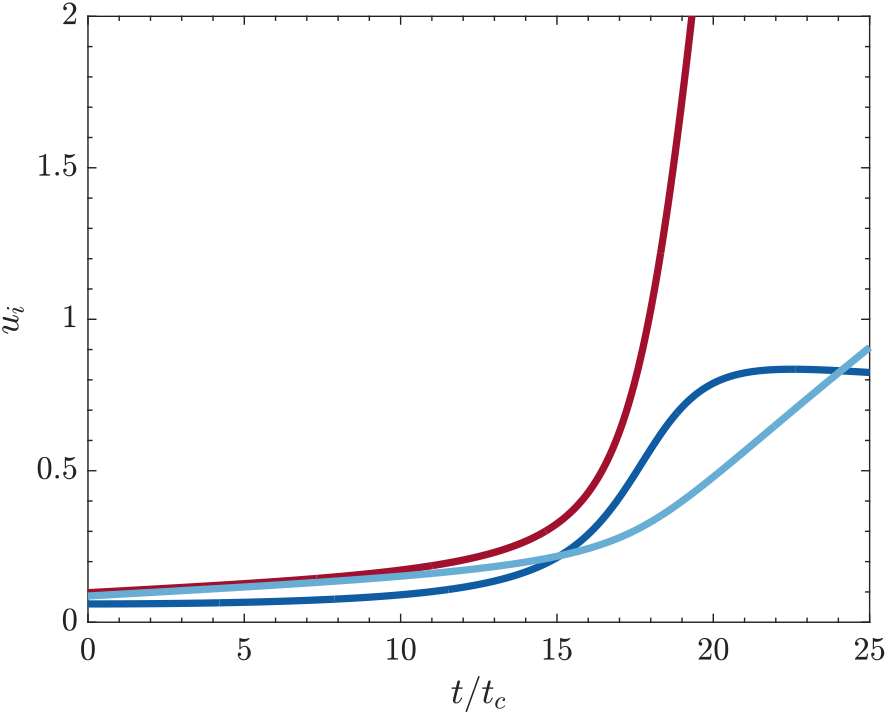
Nominal evolution of the coagulation cascade, obtained by solving eq. (6) for the 3-species coagulation model with pro-coagulant initial conditions. 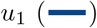, 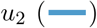 and 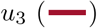

### Numerical Methods

We used the in-house code TUCAN [16, 41] to solve the Navier-Stokes equations for Newtonian, incompressible flow in the configuration described in the previous section. The numerical discretization used second-order finite differences on a Cartesian staggered grid. The temporal integration was performed with a three-stage, low-storage, semi-implicit Runge-Kutta scheme. The no-slip boundary condition at the vessel walls was modeled by the immersed boundary method [56]. After discarding initial transient effects, the velocity field (**v**(**x**, *t*)) computed by TUCAN was sampled at constant time intervals (*t*_*samp*_ = *t*_*cycle*_/35) and stored to be linearly interpolated for integrating equations (1), (4) and (14). To assess the convergence of the velocity field, we performed a grid refinement study employing three resolutions, namely, Δ*x*/*H* = 1/38, Δ*x/H* = 1/75 and Δ*x*/*H* = 1/150. Each simulation was run with a constant time step Δ*t* that ensured the Courant number to be *CFL* = max(|*u*(**x**)|)Δ*t*/Δ*x* ≈ 0.1. Richardson extrapolation was used to estimate the relative error,

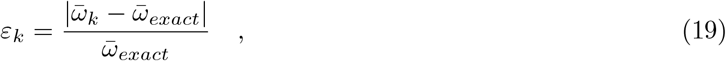

Where

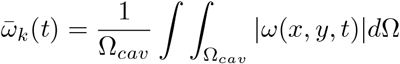

is the averaged absolute value of the vorticity in the cavity computed with a spatial resolution Δ*x* = *H/k*, Ω_*cav*_ is the volume of the cavity and 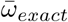 is the extrapolated value [49]. Table 2 displays the values of the relative error at three time instants (*t/t*_*c*_ = 0, *t/t*_*c*_ = 0.33, and *t/t*_*c*_ = 0.67), for the three different resolutions. The highest resolution, Δ*x*/*H* = 1/150, was selected for the present study because it yielded a relative error consistently below 2% throughout the cardiac cycle.

**Table 2:** Grid refinement study.

| Relative Error | $t/t_c = 0$ | $t/t_c = 0.33$ | $t/t_c = 0.67$ |
| --- | --- | --- | --- |
| $\varepsilon_{38}$ | 0.0904 | 0.0510 | 0.0108 |
| $\varepsilon_{75}$ | 0.0359 | 0.0185 | 0.0285 |
| $\varepsilon_{150}$ | 0.0142 | 0.0067 | 0.0108 |

A third-order weighted essentially non-oscillatory (WENO) scheme [27] was implemented to integrate the advection-reaction-diffusion PDEs (eqs. (1), (4), and (14)) as in previous works [18, 20]. This scheme locally adjusts numerical diffusivity to damp convective fluxes perpendicular to sharp scalar fronts, preventing spurious oscillations while at the same time keeping the overall numerical diffusivity low. The systems of PDEs (1), (4) and (14), and the system of ODEs (6) were integrated in time using an explicit, low-storage, 3-stage Runge Kutta scheme. In the systems of PDEs, uniform initial conditions were used for all variables, 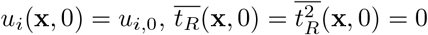, while for the ODE system the corresponding initial conditions were imposed, *u*_*i*_(0) = *u*_*i*,0_.

## Results

### Flow patterns and residence time

Pulsatile flow in the parent channel has two distinct phases coinciding with the acceleration and deceleration of the inflow profile prescribed at the inlet (Fig 2). The deceleration phase occurs for 0 ≲ *t/t*_*c*_ ≲ 0.5, whereas the acceleration phase comprises the rest of the cycle. Fig 4 shows instantaneous vorticity fields (panels A–C) and streamlines (panels D–F) at three different instants of the cycle. At the onset of deceleration (*t* = 0, first column in Fig 4), the velocity is maximum in the parent channel and a counter-clockwise vortex is the dominant pattern inside the cavity. As deceleration proceeds (*t* = 0.33*t*_*c*_, center column in Fig 4), the counter-clockwise vortex is partially sucked into the parent channel, creating a thin jet that drives fluid into the cavity in the downstream neck region while fluid slowly exits the cavity along the rest of the neck region. Finally, during acceleration (*t* = 0.67*t*_*c*_, last column in Fig 4), a vortex pair appears inside the cavity and pulls fluid from the parent channel near the upstream neck region, and ejects fluid to the channel near the downstream neck region.

**Figure 4:**
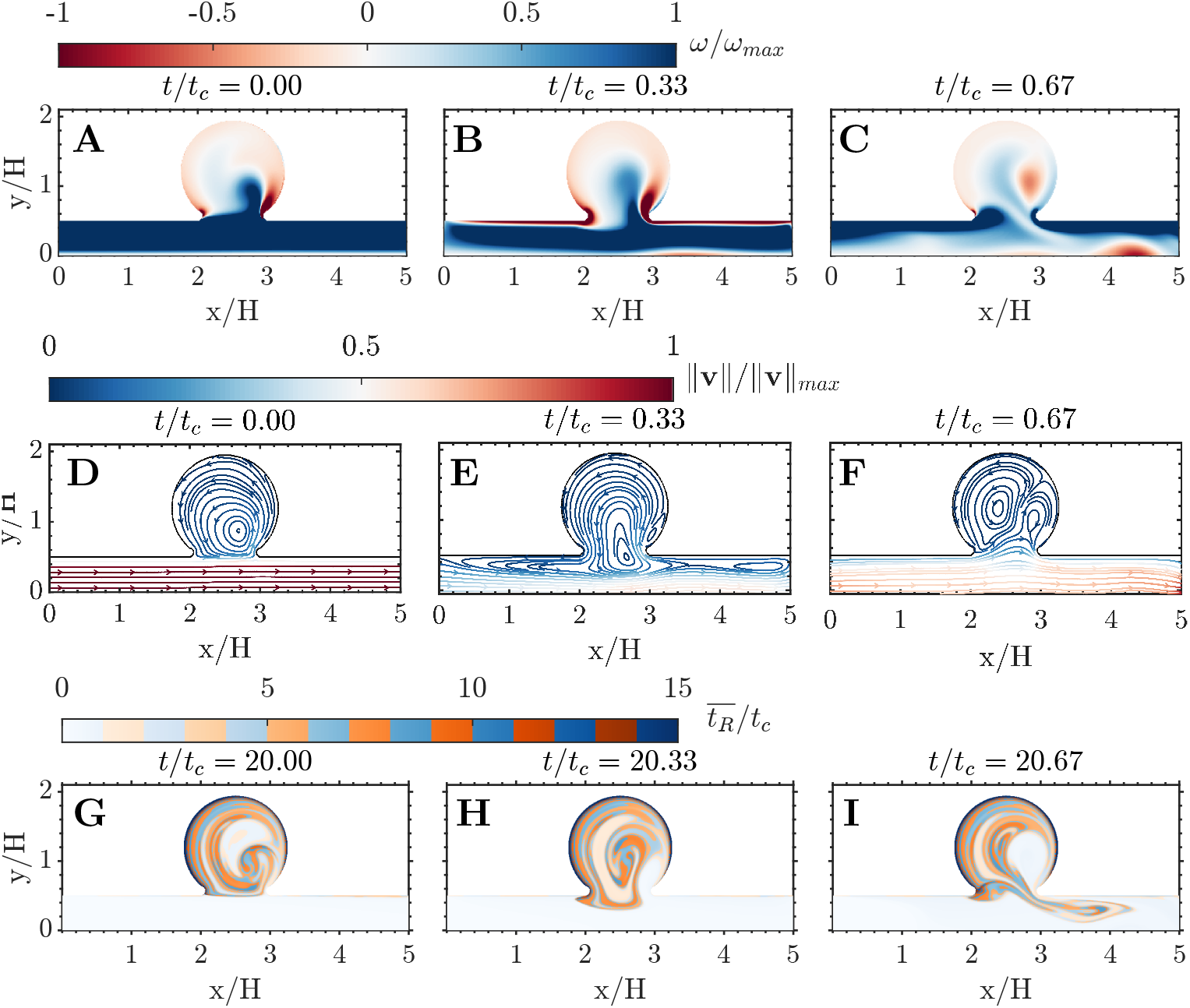
Flow patterns in the cavity. A-C: Vorticity and D-E: instantaneous streamlines colored by velocity magnitude, both normalized by its maximum value across the cardiac cycle. G-I: Residence time during the 21th simulation cycle. Each variable is plotted at three different phases of the cardiac cycle, as indicated on top of each panel.

This alternating transport of fluid into and out of the cavity repeats every cycle, generating a layered structure in residence time values reminiscent of the growth rings in a tree trunk. Figs. 4G, 4H and 4I highlight the developed layer pattern in the 21th cycle of the simulation, demonstrating that the WENO scheme effectively reproduces sharp 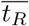 gradients over long periods of time without introducing excessive numerical diffusivity. However, albeit small, the WENO scheme does introduce a non-zero *D*_*n*_ and, consequently, this scheme leads to *σ*_*T*_ > 0. This can be observed in Fig. 5, which displays snapshots of 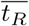 and *σ*_*T*_ over the period 10*t*_*c*_ ≤ *t* ≤ 21*t*_*c*_. Both variables grow with time, while keeping approximately the same spatial organization.

**Figure 5:**
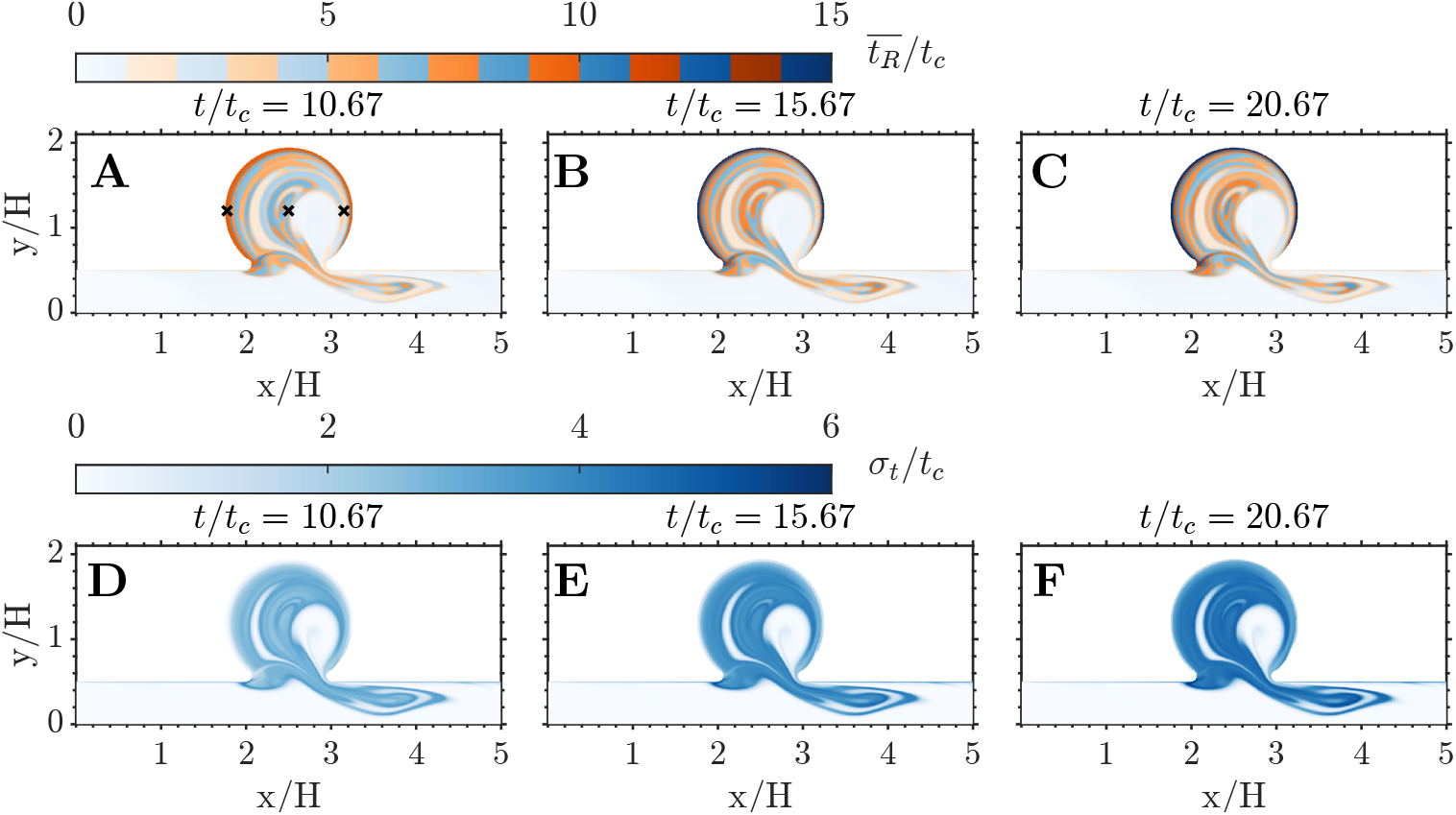
Mean and standard deviation of the residence time. A-C: spatial distribution of 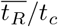. D-F: spatial distribution of *σ*_*T*_ */t*_*c*_. Each variable is plotted at three different cycles. A,D: 11th cycle. B,E: 16th cycle. C,F: 21th cycle.

Fig. 6 presents the temporal evolution of 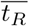 and *σ*_*T*_ at three points along the horizontal diameter of the cavity (i.e., the crosses in Fig. 5A). In all three points, the residence time exhibits an initial phase of linear growth, 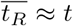 and *σ*_*T*_ ≈ 0, as in our previous work [18]. At points that do not receive “fresh” fluid from the parent channel (e.g., the wall point in Fig. 6A), this phase should last indefinitely. In our simulations, a small departure from 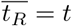 and *σ*_*T*_ = 0 becomes noticeable for *t* ≳ 15*t*_*c*_ due to the WENO scheme’s numerical diffusivity. Points exchanging fluid with the parent channel experience a different behavior characterized by two principal features. First, 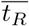 rises and falls every cardiac cycle as pockets of stagnant and fresh fluid move back and forth over the point of interest. Second, the envelope of 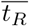 saturates to a maximum value 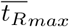 indicating the time needed for fluid exchange with the parent channel to wash out the local blood pool completely. At these points, *σ*_*T*_ follows a similar behavior since fresh blood from the parent channel has not only low 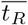 but also low *σ*_*T*_. We note that, for long enough simulation times, *σ*_*T*_ can grow at certain points to be comparable in magnitude to 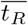. This result has implications for multi-fidelity modeling of the coagulation cascade but also for the uncertainty of 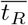 itself.

**Figure 6:**
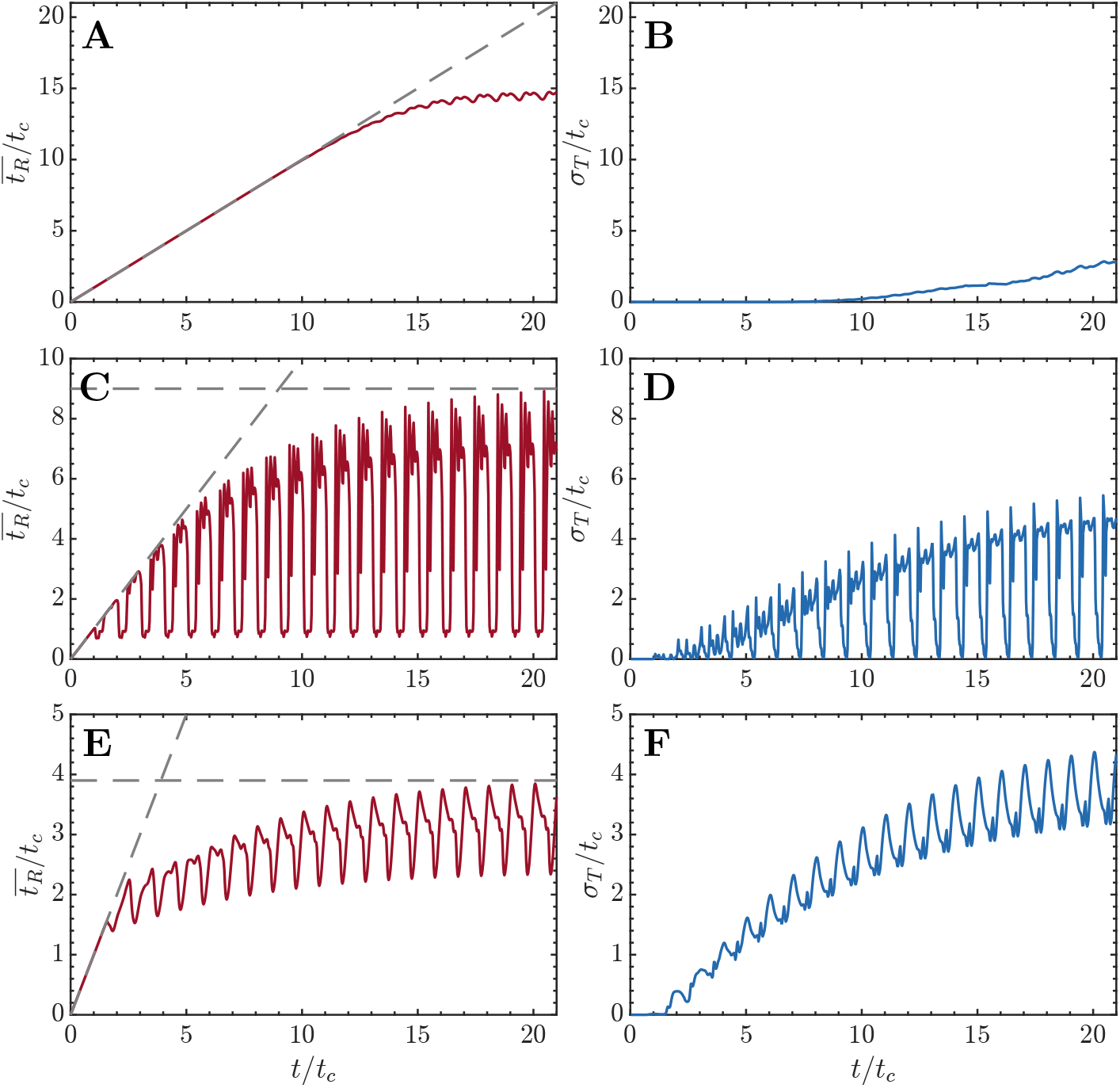
Time series of residence time and its standard deviation. A,C,E: Temporal evolution of 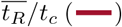. B,D,F: Temporal evolution of 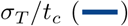. Three locations are considered, indicated with × in Fig. 5A: A,B at (*x/H, y/H*) = (1.78, 1.19); C,D at (*x/H, y/H*) = (2.5, 1.19); and E,F at (*x/H, y/H*) = (3.15, 1.19).

### Multi-Fidelity Modeling of the Coagulation Cascade

The initiation of coagulation is characterized by a rapid growth in the concentration of the three species, especially thrombin (*u*_1_), after 10-15 cycles (see Fig. 3). Fig. 7 depicts the spatio-temporal structure of *u*_1_ over the 16th simulation cycle, as obtained by the MuFi-1 (panels a—c) and MuFi-2 (panels d—f) models, as well as by the reference HiFi model (panels g—i). The three models capture the rise of *u*_1_ inside the cavity and the formation of a layered structure similar to that of the residence time. There is a trend for MuFi-1 to underestimate *u*_1_, which is for the most part corrected by MuFi-2.

**Figure 7:**
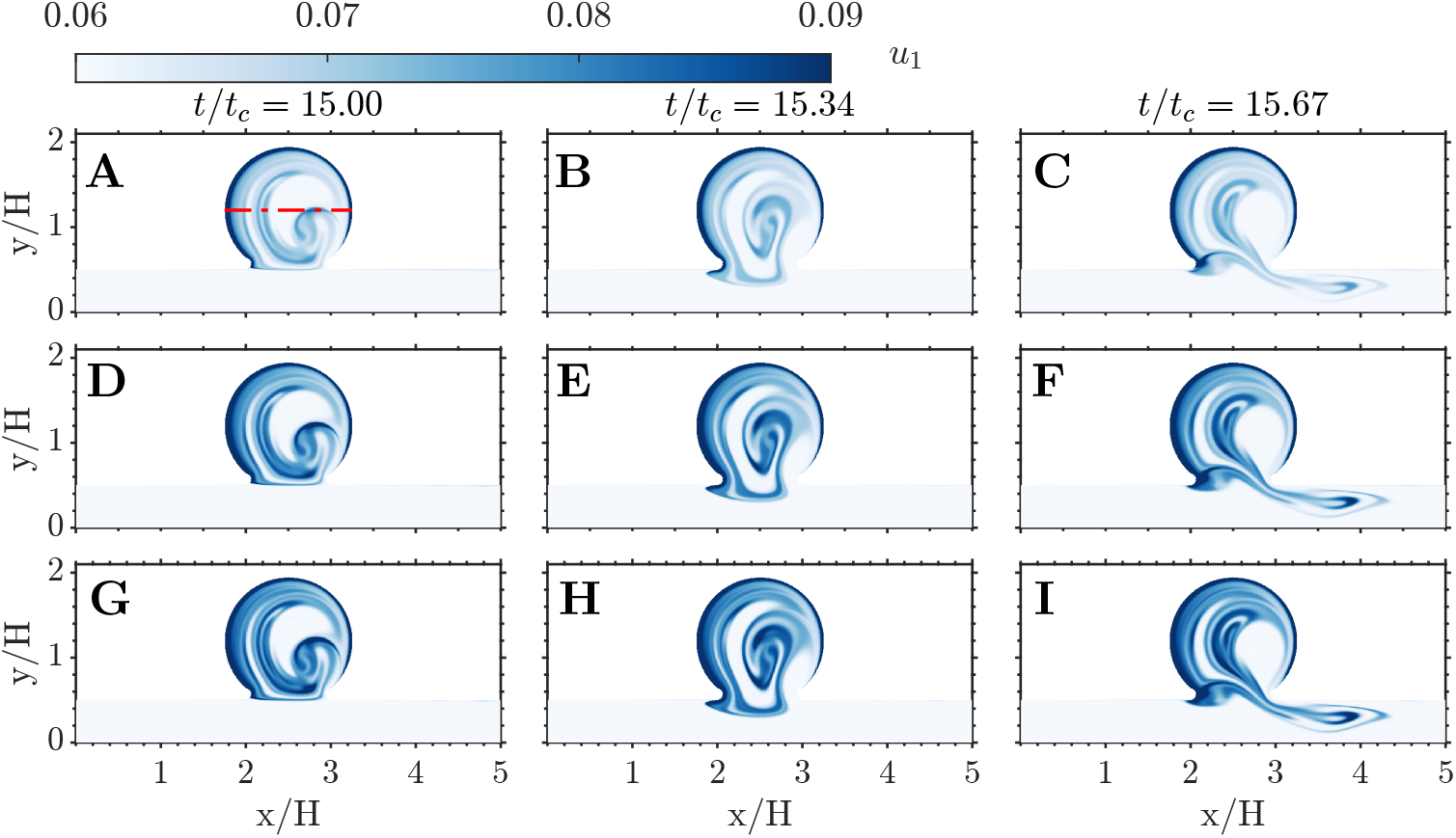
Spatial distribution of thrombin concentration. A-C: MuFi-1. D-F: MuFi-2. G-I: HiFi reference model. Three phases within the 16th cycle are plotted for each case, as indicated on the top row. A,D,G: *t/t*_*c*_ = 15. B,E,H: *t/t*_*c*_ = 15.34. C,F,I: *t/t*_*c*_ = 15.67.

To study each model’s behavior in more detail, Fig. 8 shows the thrombin concentration vs. time at the same three points considered in Fig. 6, representative of scenarios where 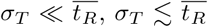, or 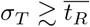 after running the simulation for a large number of cycles, 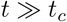. For reference, the figure shows also the thrombin concentration obtained directly from the HiFi models and the ODE eqs. (6), the latter corresponding to no-flow and zero diffusion, i.e., 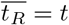 and *σ*_*T*_ = 0.

**Figure 8:**
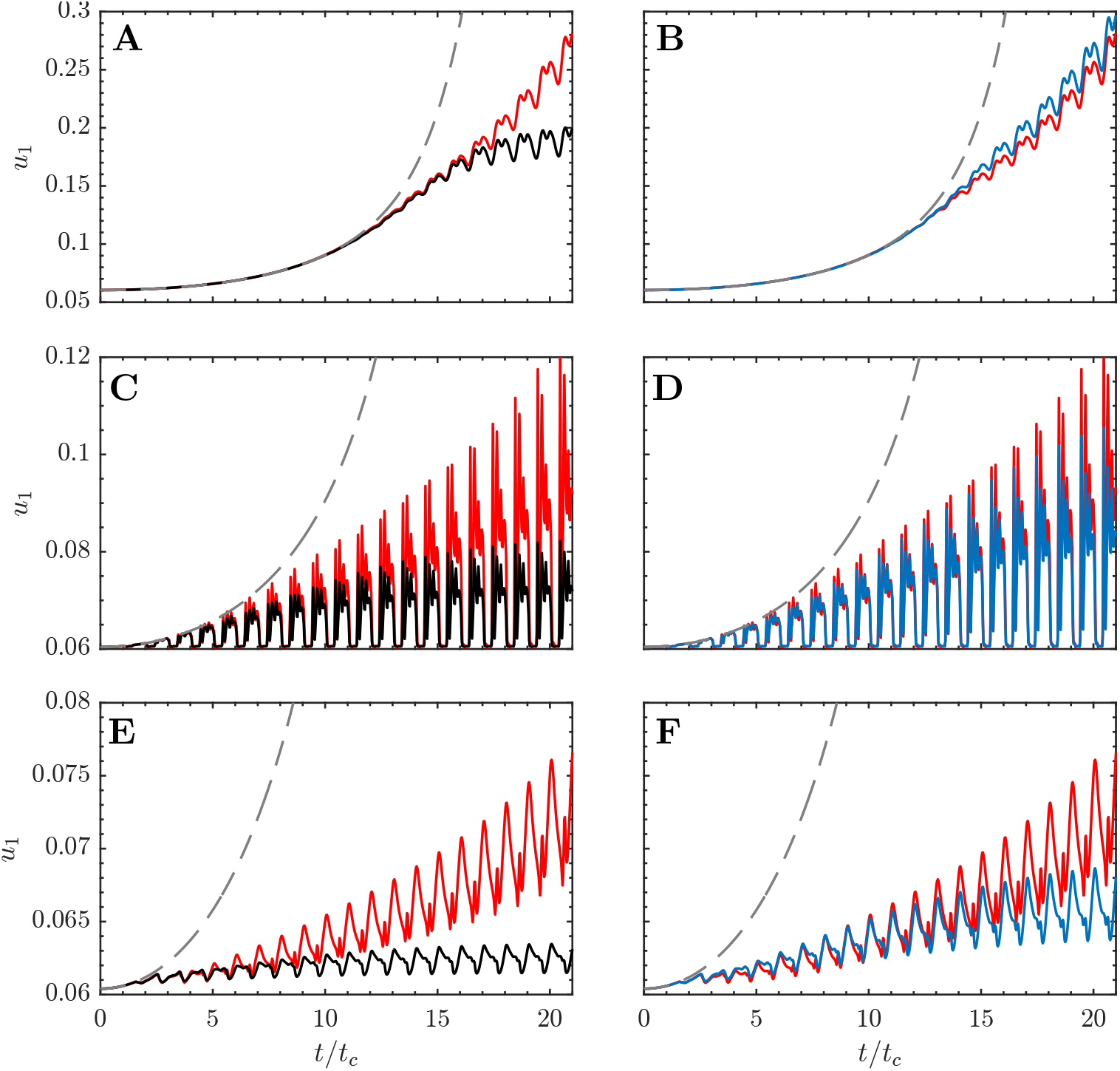
Time series of thrombin concentration, *u*_1_. Each line correspons to a different model: MuFi-1 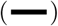, MuFi-2 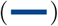 and HiFi model 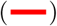. For reference, the solution of the 3-ODE system eq. (6) is also included 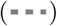. Three locations are considered, indicated with × in Fig. 5A: A,B at (*x/H, y/H*) = (1.78, 1.19); C,D at (*x/H, y/H*) = (2.5, 1.19); and E,F at (*x/H, y/H*) = (3.15, 1.19).

Similar to the residence time, *u*_1_ experiences peaks and valleys every cycle due to the periodic fluid exchange between the cavity and the parent channel. In addition, the peak value of *u*_1_ increases from one cycle to the next as the coagulation cascade progresses in the fluid trapped in the cavity. The zero-flow ODE model (dashed lines in Fig. 8) fails to capture the oscillations of *u*_1_ and severely overestimates the growth of its peak values. The MuFi-1 model captures the oscillatory nature of *u*_1_, but it begins to underpredict the growth of the peak values after a number of cardiac cycles that varies from point to point. At the first sampled point, where 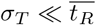, the MuFi-1 model remains accurate for *t* ∼ 15*t*_*c*_ (figure 8a), while at the second and third sampled points, where 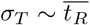, the MuFi-1 model significantly departs from the HiFi model beyond *t* ∼ 10*t*_*c*_ (Fig. 8C, E). The MuFi-2 model, which incorporates both 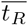 and *σ*_*T*_, remains in reasonable agreement with the HiFi model much longer than the MuFi-1 model, producing fairly accurate results for simulation times well over 10 cycles at the three sampled points (Fig. 8B, D, F). By *t* = 20*t*_*c*_, the MuFi-1 model underestimates the peak *u*_1_ by 22%, 33.8%, and 16% while the MuFi-2 model does so by 6%, 15.3%, and 8.9%, respectively, at the first, second, and third sampled points.

We compare the spatio-temporal behavior of the three species across the high-fidelity and multi-fidelity models by representing in Fig. 9 the concentrations *u*_1_, *u*_2_, and *u*_3_ along the horizontal cavity diameter depicted in Fig. 7A, with *y/H* = 1.19. Spatial concentration profiles are plotted at time-points (*t/t*_*c*_ = 10, 15, 20). Factor XIa (*u*_2_) displays the best agreement among models, followed by the inhibitor PCa (*u*_3_) and thrombin (*u*_1_). Of note, the MuFi-2 model accurately replicates the HiFi behavior up to *t* ≈ 15*t*_*c*_ for all three species, capturing the spatial complexity of the HiFi oscillations. In comparison, the accuracy of the MuFi-1 model deteriorates faster with simulation time, although this model still retains the qualitative spatial dependence of species concentration.

**Figure 9:**
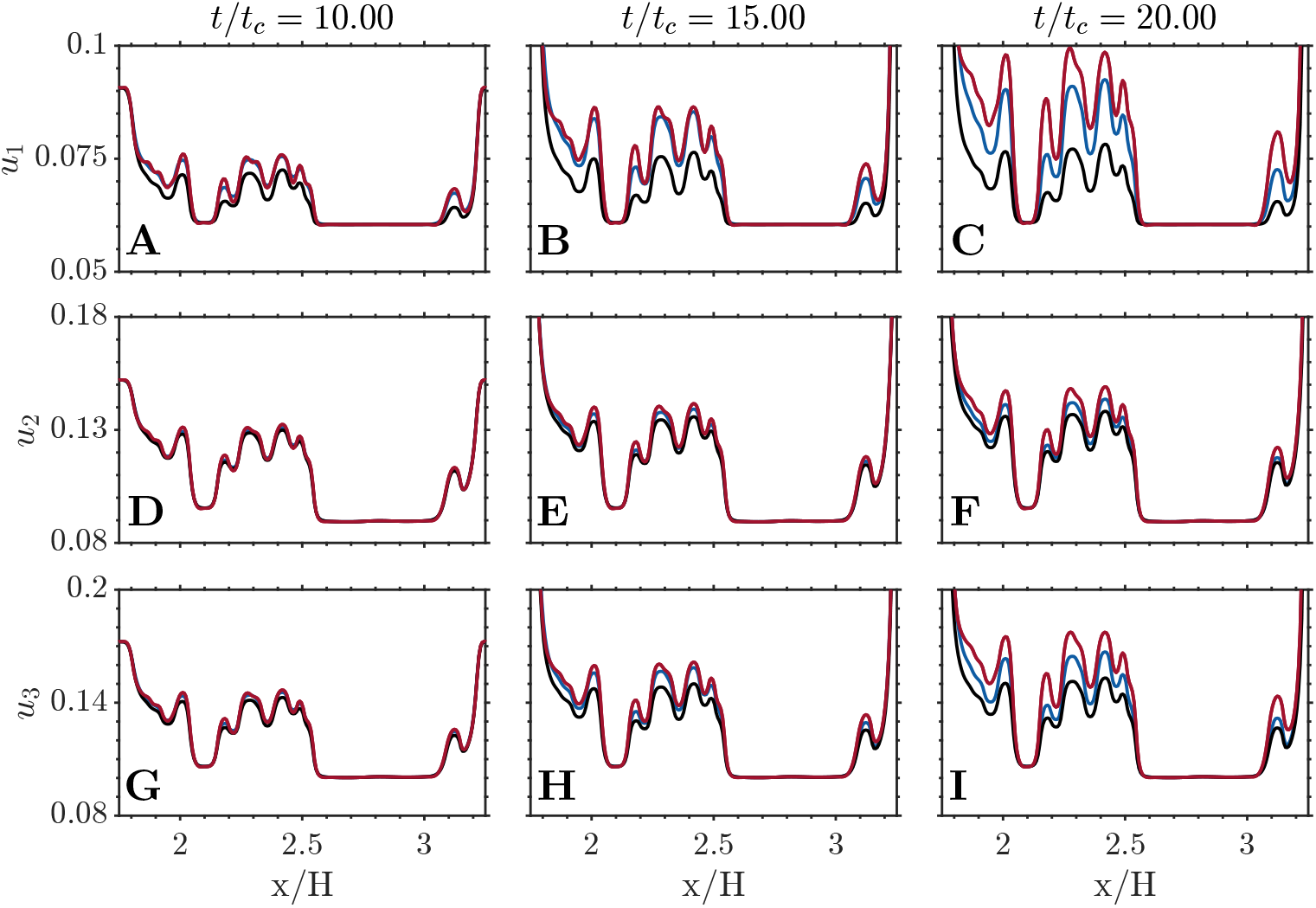
Spatial distribution of species concentration. Data is show at *y/H* = 1.19 for MuFi-1 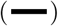, MuFi-2 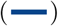 and HiFi model 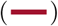. A,B,C: thrombin, *u*_1_. D,E,F: factor XIa, *u*_2_. G,H,I: protein C activated, *u*_3_. Three different times are plotted for each species, as indicated on the top row.

### Multi-Fidelity Model Error Analysis

While interesting to understand the performance of the multi-fidelity models, Figs. 8 and 9 include examples of extreme behavior that do not reflect these models’ overall accuracy. To systematically quantify the discrepancies between the high-fidelity and multi-fidelity models, we compute the relative errors

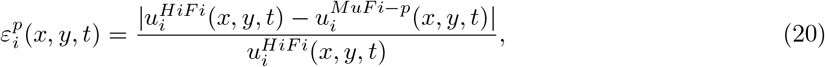

where *i* stands for species and *p* for MuFi order. Fig. 10 shows the thrombin relative errors of the MuFi-1 and MuFi-2 models, 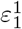 and 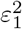, for three instants along the 16th simulation cycle. These errors are negligible in the parent channel but reach appreciable values inside the cavity, where they exhibit a layered pattern with strong gradients, similar to 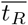 and *σ*_*T*_. Also, 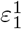 is significantly higher than 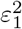, consonant with the data shown in Figs. 7–9.

**Figure 10:**
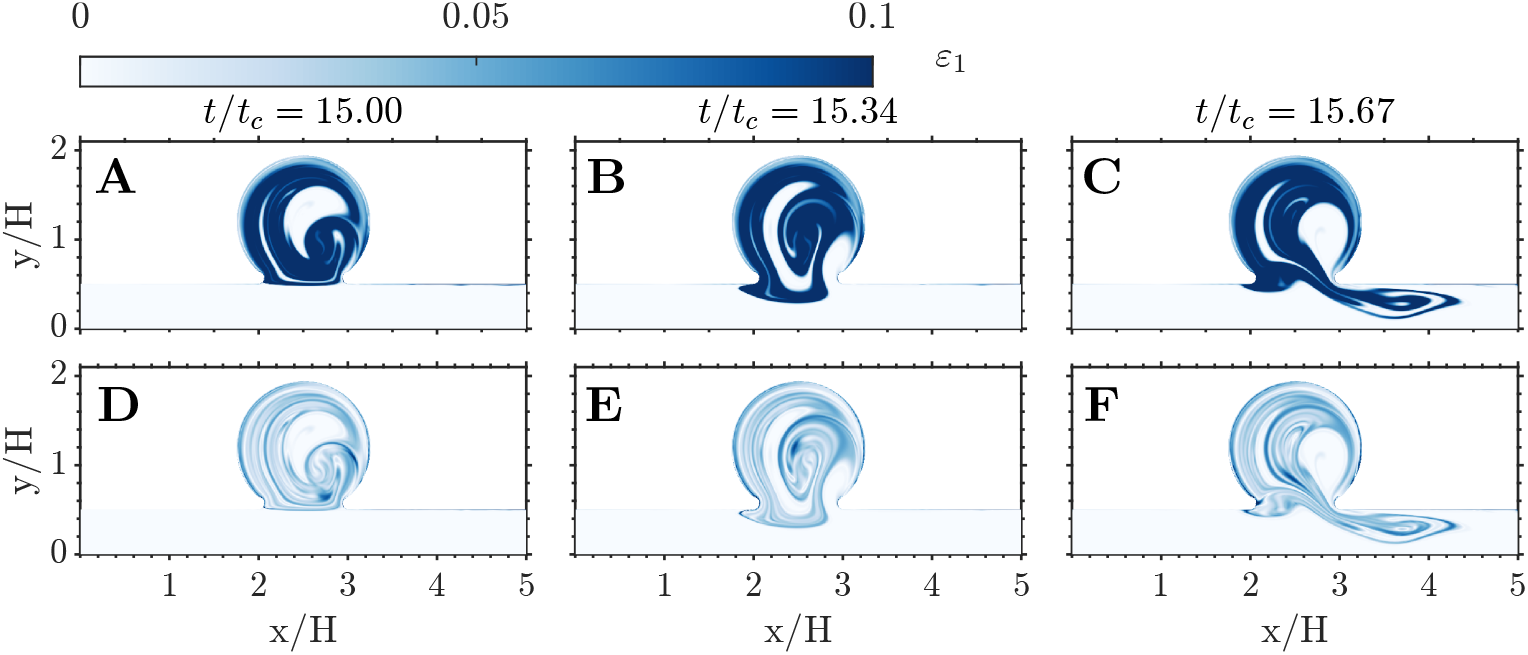
Spatial distribution of relative errors in thrombin concentration. The relative error 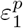 is defined in eq. (20). A-C: MuFi-1. D-F: MuFi-2. Three phases within the 16th cycle are plotted for each case, as indicated on the top row. A,D,G: *t/t*_*c*_ = 15. B,E,H: *t/t*_*c*_ = 15.34. C,F,I: *t/t*_*c*_ = 15.67.

To characterize the dependence of the MuFi errors on the residence time and its variance inside the cavity, we divide the 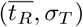 plane in bins, ensemble-average 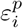 inside each bin, and plot the resulting error maps in Fig. 11 together with the corresponding normalized bin counts [i.e., the probability density function 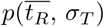]. The error maps obtained for different values of *t/t*_*c*_ indicate that the MuFi error increases with *σ*_*T*_ while it is much less sensitive to 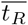. The MuFi-2 model particularly outperforms the MuFi-1 model in areas of large *σ*_*T*_, since MuFi-1 assumes *σ*_*T*_ = 0. Similar observations can be made for the error maps for the concentrations of factor XI and PCa, which are provided in S4 Appendix. Inspection of the probability density functions shows that the majority of the points inside the cavity are circumscribed to two regions in the 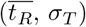 plane. The region near the point 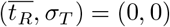 corresponds to locations within the cavity that receive periodic inflows of fresh flow from the parent channel. This periodic infusion sustains a low level of 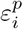 throughout each cycle, since fresh flow has small values of 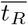 and 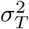. The region near 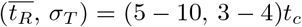 corresponds to the stagnant areas of recirculating flow inside the cavity(see Fig. 5). The modal errors in this region (black crosses in Fig. 11G–I) are 0.032, 0.096, and 0.2 for MuFi-1 and 0.007, 0.015, and 0.067 for MuFi-2 at *t/t*_*c*_ = 9.5, 14.5, and 19.5, respectively. These stagnant flow areas displace upwards and towards the right in the 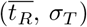 plane as the simulation time advances (see bottom row of Fig. 11). Consequently, we expect *σ*_*T*_ and 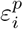 to grow with *t/t*_*c*_ inside the cavity and the MuFi-2 model to outperform the MuFi-1 model.

**Figure 11:**
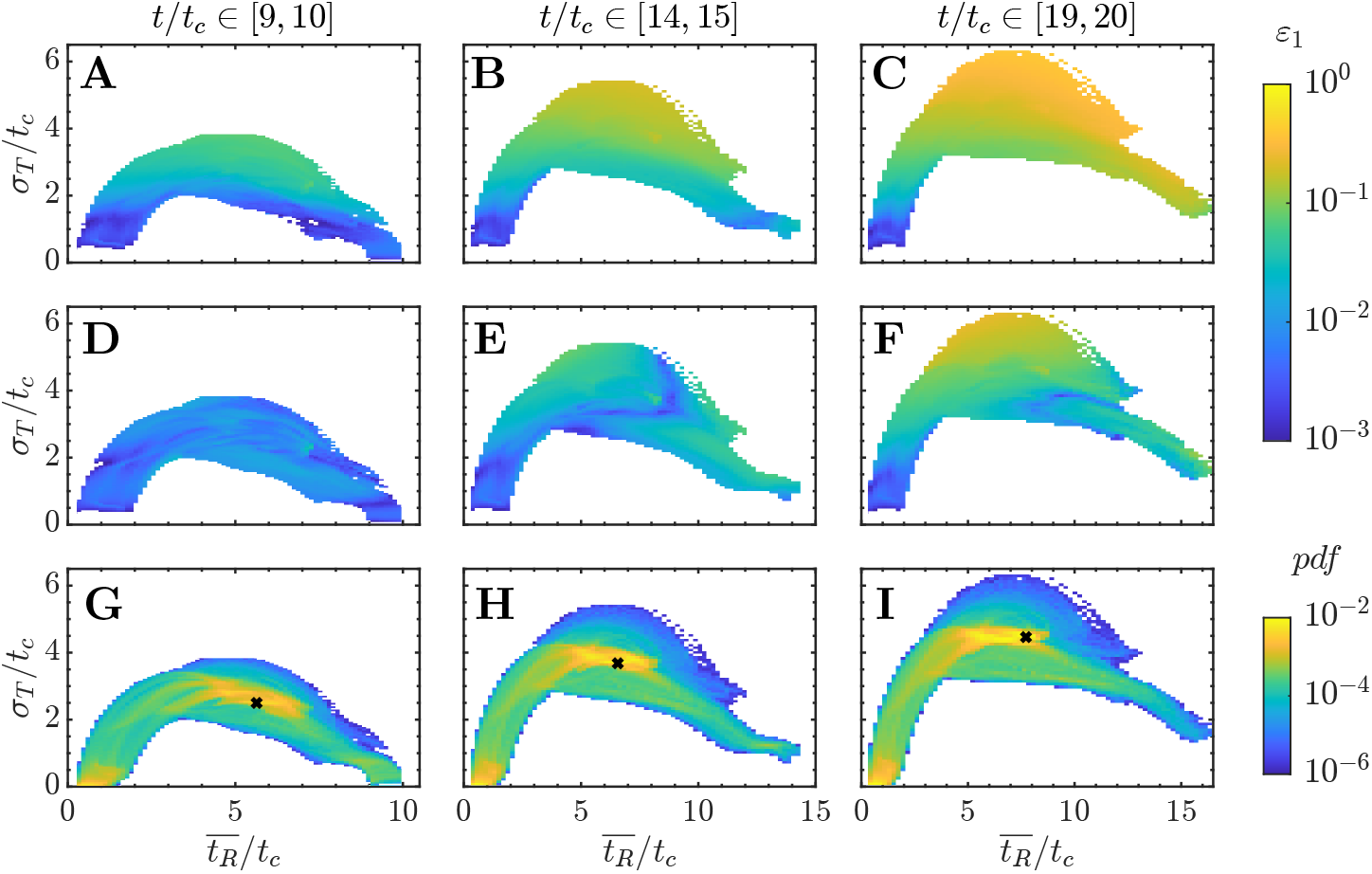
Error maps for thrombin. A-C: Relative error in thrombin concentration in MuFi-1, 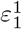, as a function of residence time and its standard deviation. D-F: Same, but for MuFi-2, 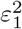. G-I: Joint probability density function of residence time 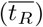 and its standard deviation (*σ*_*T*_). Data for all panels is compiled inside cavity during three different cycles, as indicated on the top row. A,D,G: 10th cycle. B,E,H: 15th cycle. C,F,I: 20th cycle.

To test this hypothesis and quantify the overall error of MuFi models, we calculate the spatial average of 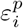 over the cavity region,

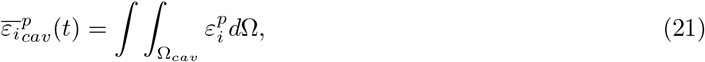

and plot it vs. simulation time in Fig. 12 for the three species. At short times (*t*, :S 4*t*_*c*_), the MuFi-1 and MuFi-2 models produce similar errors with respect to the HiFi model, which scale approximately as (*t/t*_*c*_)^3^. These errors average to less than 0.5% (i.e., 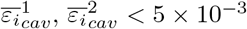) and are markedly lower for thrombin than for the other two species. For simulation times (4*l*_*c*_ ≳ *l* ≲ 10*l*_*c*_), the MuFi-1 model’s error continues to grow approximately as (*t/t*_*c*_)^3^ for all three species while the MuFi-2 error almost plateaus. Thus, at *t* = 10*t*_*c*_, MuFi-1 model remains generally acceptable (errors below 4%), while the MuFi-2 model still provides excellent accuracy (errors below 0.6%). However, beyond 10*t*_*c*_, the relative errors for all three scalars in the MuFi-2 model begin to grow again roughly as (*t/t*_*c*_)^3^. By *t* = 20*t*_*c*_, the MuFi-2 model offers an acceptable 5% global error 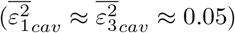 but the MuFi-1 errors can grow to almost 15% 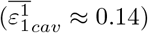.

**Figure 12:**
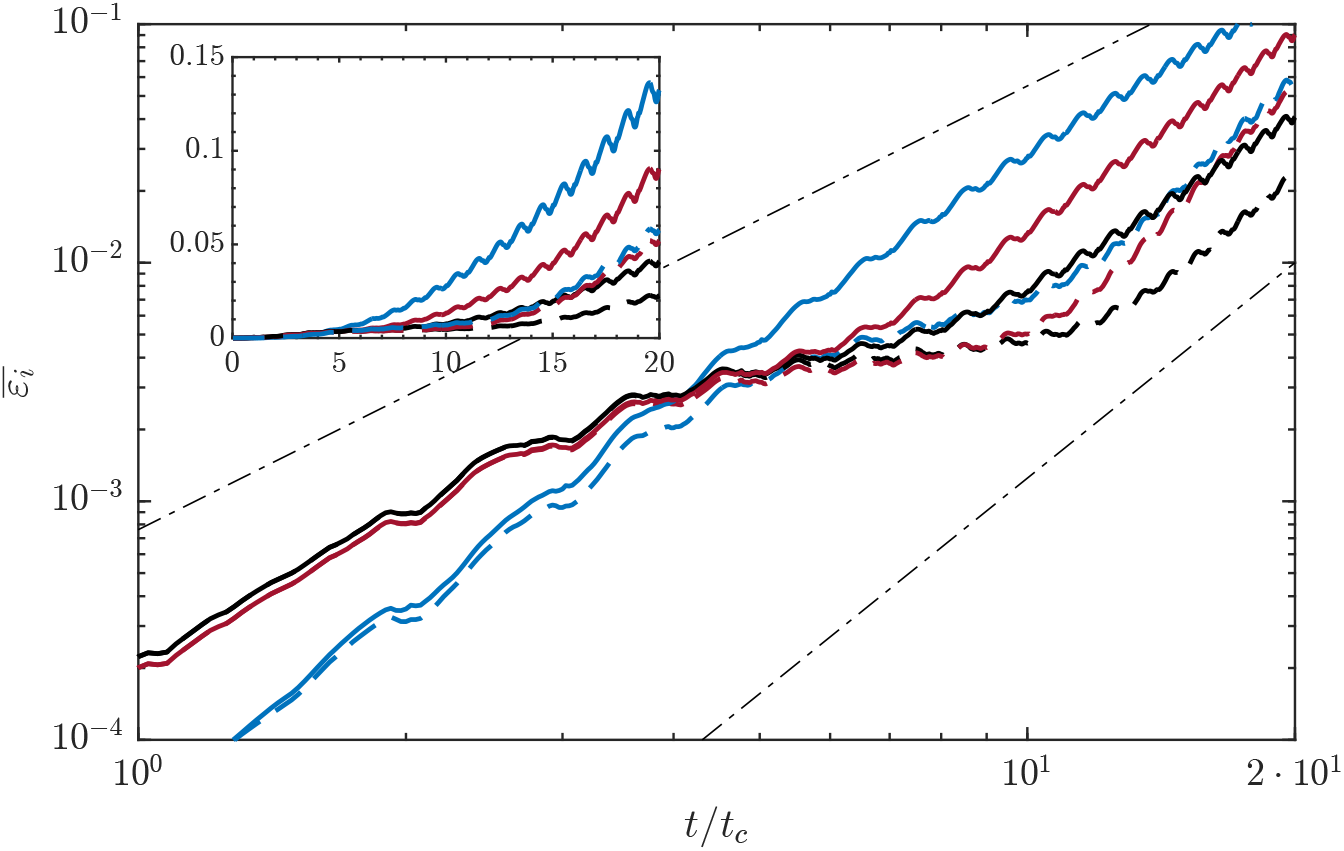
Averaged relative error in the cavity for MuFi-1 (solid) and MuFi-2 (dashed), 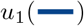, 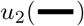 and 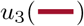, dashed-dot lines for 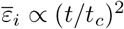 and 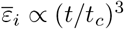, top and bottom respectively.

## Discussion

The computational modeling of the coagulation cascade in flowing blood poses significant challenges due to the multi-scale nature of the process and the large number of involved chemical species. High-fidelity (HiFi) continuum mechanics models of coagulation lead to dozens of reaction-advection-diffusion partial differential equations (PDEs) [36]. The reaction kinetics in these equations are much slower than the cardiac cycle, requiring long simulation times to cover the coagulation process [4, 28, 38]. Moreover, the diffusion of chemical species is orders of magnitude slower than their convection and reaction kinetics, causing sharp scalar fronts that require ultra-high-resolution spatial meshes [29]. Despite the vast advances in computational software and hardware achieved in past decades, these joint requirements still impede the fast simulations of chemo-fluidic coagulation models in arterial geometries, hindering the adoption of these models in clinical decision-making. Modelers usually resort to surrogate metrics associated with thrombogenesis, derived from blood residence time or wall shear stress [3, 12, 31, 43, 48, 51]. On the other hand, chemo-fluidic models of thrombosis are rare [6, 53] and often suffer from excessive diffusivity (numerical or explicit), short simulation times, and/or simplified coagulation models with few species.

We introduce a family of tailorable multi-fidelity (MuFi) models to reduce the computational cost of simulating the coagulation cascade of *N* species in a flow. The MuFi models are designed to approximate the HiFi model in the limit of vanishing molecular diffusivity of the reacting species. In this limit, the *N* -PDEs of the HiFi model can be transformed into ordinary differential equations (ODEs) by changing variables between simulation time and blood residence time. Consequently, the HiFi model is replaced by *N* ODEs representing the reaction kinetics and *p* PDEs representing the ensemble mean residence time within each fluid particle, 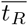, and *p* − 1 higher-order statistical moments (namely, 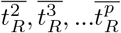) [11]. We provide a procedure to incrementally derive MuFi models of arbitrary order starting from the first-order model obtained with *p* = 1, corresponding to zero diffusivity. In the presence of small diffusivity, natural or numerical, higher-order models can be derived by Taylor-expanding the HiFi model around the zero diffusivity limit.

We assess the performance of the MuFi-1 and MuFi-2 models obtained for *p* = 1, 2 in a well-characterized, simplified coagulation system representing three species: thrombin, activated factor XI, and activated protein C [62]. This coagulation system is evaluated in a pulsatile flow through a two-dimensional geometry consisting of a parent channel driven by a Womersley inflow profile with a laterally protruding cavity where blood becomes stagnant. The non-dimensional parameters governing the flow, i.e., the Reynolds (*Re* = 500) and Womersley (*α* = 10) numbers, are representative of a saccular aneurysm in an intermediate-size artery of the adult human circulatory system [58]. For the residence time and its variance, the Peclet number is nominally set to be infinitely high by not including mass diffusivity terms in the corresponding PDE equations.

This configuration creates a cyclic influx of fresh fluid from the parent channel to the cavity and efflux of stagnant fluid from the cavity to the channel, together with a swaying fluid motion inside the cavity. Flow transport leads to a complex residence time pattern with thin layers separated by strong gradients, which intensifies as 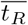 grows with simulation time. Our WENO scheme resolve the thinly layered 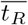 pattern for long simulation times. However, albeit low, the WENO scheme has a non-zero numerical diffusivity [27], and thus, the effective *Pe* in our HiFi simulations is finite, affecting the solution to the transport equations for 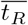 and 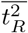. Consequently, not only 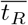 but also its standard deviation, *σ*_*T*_, increases with simulation time.

Overall, the MuFi-1 model compares well with the HiFi model, producing spatially averaged errors inside the cavity that remain below 5% for up to 10 simulation cycles even if this model was derived under the assumption of *Pe* →∞ and *σ*_*T*_ = 0. Nevertheless, MuFi-1 starts to underestimate the concentration of all species beyond that point. By *t* = 20*t*_*c*_, the spatially averaged errors inside the cavity are as high as ≈ 15%. The MuFi-2 model yields significantly lower errors than the MuFi-1 model, with spatially averaged errors that remain below 5% for up to *t* ≈ 20*t*_*c*_. At short simulation times, the MuFi models are most accurate for thrombin, but this behavior is reversed for *t* .2’ 4*t*_*c*_. Statistical analysis shows that the MuFi errors depend almost exclusively on *σ*_*T*_ and that this dependence is steeper for MuFi-1 than for MuFi-2. Because *σ*_*T*_ increases with numerical diffusivity, *D*_*n*_, the performance of MuFi implementations is expected to depend on the numerical scheme and the spatio-temporal resolution used to discretize the model PDEs. While an exhaustive analysis of the numerical diffusivity of numerical discretizations of advection-reaction-diffusion problems is beyond the scope of this study, we have outlined a general methodology to formulate MuFi models of arbitrary order, derive each numerical scheme’s EDE for *σ*_*T*_, and outlined a step-by-step process to compute *σ*_*T*_ without the need to derive its EDE. With these tools, it should be straightforward to tailor MuFi models to the peculiarities of each flow and numerical solver. These step-by-step procedures should apply to models that compute residence time using non-classical numerical representations of the governing PDEs such as, e.g., neural networks.

The MuFi models (*N* ODEs, *p* PDEs) are much more efficient than the HiFi model (*N* PDEs) because solving ODEs is significantly cheaper than solving PDEs. In physiologically relevant computational meshes containing hundreds of elements in each direction, MuFi models achieve a speedup > *N/p*. We anticipate this speedup to exceed an order of magnitude, considering the favorable accuracy achieved by low-order MuFi models (*p* = 1, 2) and the dozens of species involved in realistic coagulation cascade models. An additional advantage of MuFi models is that one can run any number of different models inexpensively for a specified flow baseline. The costly part of MuFi models, i.e., solving the p-PDEs representing residence time and its higher-order moments, does not need to be redone unless the anatomy or inflow/outflow conditions change. Thus, MuFi models are highly efficient for sensitivity analyses, uncertainty quantification, kinetics model comparisons, and any type of study requiring multiple evaluations of the coagulation cascade model. For simplicity and to establish proof of concept of the MuFi strategy, this study focuses on the intrinsic coagulation cascade, ignoring the extrinsic cascade initiated by the release of procoagulatory or inhibitory species from the vessel walls. While both the intrinsic and extrinsic coagulation pathways are often activated in cardiovascular conditions associated with thrombosis [7, 8, 13], focusing on the intrinsic pathway is not uncommon in the literature, motivated by the paucity of information about the wall’s prothrombotic potential in many clinically relevant scenarios [21]. Generating MuFi models for the extrinsic pathway would involve additional PDEs representing the residence time of wall-released chemical species or the time spent by blood near damaged wall regions, e.g., the near-wall residence time proposed by others [53]. As long as the number of additional PDEs remains lower than the total number of species, *N*, the resulting

MuFi models would still save computational time. Finally, note that residence time and its higher order moments can not only be obtained *in silico* using CFD analysis [18,-20,-48], but also *in vitro* using experimental techniques like particle image velocimetry [15, 40, 55], and *in vivo* using medical imaging modalities like phase-contrast magnetic resonance imaging or Doppler ultrasound [23, 37, 54]. Residence time maps obtained *in vitro* or *in vivo* could be fed to the ODE component of the MuFi models, providing a computationally tractable method to calculate the spatio-temporal evolution of the coagulation species from multiple data sources. In particular, this strategy would offer promise for non-invasively imaging thrombogenesis in the clinical setting.

## Conclusion

We present a novel multi-fidelity approach to mitigate the computational burden of simulating the coagulation cascade under flow. Using residence time 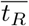 and its statistical moments, the multi-fidelity (MuFi) modeling achieves a favorable trade-off between computational cost and accuracy. The multi-fidelity approach allows for the integration of various data streams, including CFD analysis, *in vitro* experiments, and non-invasive imaging *in vivo* techniques, enabling a comprehensive understanding of the spatial and temporal progression of coagulation species and offering promise for clinical translation. Finally, we note that the same multi-fidelity strategy developed here can be adopted to increase the efficiency of simulating other systems biology processes influenced by blood flow.

## Acknowledgments

This work was partially supported by Comunidad de Madrid (Synergy Grant Y2018/BIO-4858 PREFI-CM), Spanish Research Agency(AEI, grant number PID2019-107279RB-I00), Instituto de Salud Carlos III (grant numbers PI15/02211-ISBITAMI and DTS/1900063-ISBIFLOW), the EU—European Regional Development Fund, and the US National Institutes of Health (grant numbers 1R01HL160024 and 1R01HL158667).

## References

1. N. M. Al-Saady, O. A. Obel, and A. J. Camm. Left atrial appendage: structure, function, and role in thromboembolism. Heart, 82(5):547–554, 1999.

2. M Anand, K Rajagopal, and KR Rajagopal. A model for the formation, growth, and lysis of clots in quiescent plasma. a comparison between the effects of antithrombin iii deficiency and protein c deficiency. J. Theor. Biol., 253(4):725–738, 2008.

3. Amirhossein Arzani, Ga-Young Suh, Ronald L Dalman, and Shawn C Shadden. A longitudinal comparison of hemodynamics and intraluminal thrombus deposition in abdominal aortic aneurysms. Am. J. Physiol. Heart Circ. Physiol., 307(12):H1786–H1795, 2014.

4. F. I. Ataullakhanov, Y. V. Krasotkina, V. I. Sarbash, R. I. Volkova, E. I. Sinauridse, and A. Y. Kondratovich. Spatio-temporal dynamics of blood coagulation and pattern formation: an experimental study. Int. J. Bifurcat. Chaos, 12(09):1969–1983, 2002.

5. Fazoil I Ataullakhanov, Veronika I Zarnitsyna, Andrei Yu Kondratovich, Ekaterina S Lobanova, and Vasilii I Sarbash. A new class of stopping self-sustained waves: a factor determining the spatial dynamics of blood coagulation. Phys-Usp+, 45(6):619, 2002.

6. J. Biasetti, P. G. Spazzini, J. Swedenborg, and T. C. Gasser. An integrated fluid-chemical model toward modeling the formation of intra-luminal thrombus in abdominal aortic aneurysms. Front. Physiol., 3:266, 2012.

7. A. Camaj, V. Fuster, G. Giustino, S. W. Bienstock, D. Sternheim, R. Mehran, G. D. Dangas, A. Kini, S. K. Sharma, J. Halperin, et al. Left ventricular thrombus following acute myocardial infarction: Jacc state-of-the-art review. J. Am. Coll. Cardiol., 79(10):1010–1022, 2022.

8. S. J. Cameron, H. M. Russell, and A. P. Owens III. Antithrombotic therapy in abdominal aortic aneurysm: beneficial or detrimental? Am. J. Hematol., 132(25):2619–2628, 2018.

9. J. E. Cohen, E. Yitshayek, J. M. Gomori, S. Grigoriadis, G. Raphaeli, S. Spektor, and G. Rajz. Spontaneous thrombosis of cerebral aneurysms presenting with ischemic stroke. J. Neurol. Sci, 254(1-2):95–98, 2007.

10. Earl W Davie and Oscar D Ratnoff. Waterfall sequence for intrinsic blood clotting. Science, 145(3638):1310–1312, 1964.

11. J. C. del Alamo, L. Rossini, A. Kahn, J. Bermejo, P. Martínez-Legazpi, and R. Y. Alvarez. Mapping and quantifying blood stasis and thrombus risk in the heart, July 21 2020. US Patent 10,716,519.

12. P Di Achille, G Tellides, CA Figueroa, and JD Humphrey. A haemodynamic predictor of intraluminal thrombus formation in abdominal aortic aneurysms. Proc. Math. Phys. Eng. Sci., 470(2172):20140163, 2014.

13. W. Y. Ding, D. Gupta, and G. Y. H. Lip. Atrial fibrillation and the prothrombotic state: revisiting virchow’s triad in 2020. Heart, 106(19):1463–1468, 2020.

14. E. A. Ermakova, M. A. Panteleev, and E. E. Shnol. Blood coagulation and propagation of autowaves in flow. Pathophysiol. haemost. thromb., 34(2-3):135–142, 2005.

15. A. Falahatpisheh and A. Kheradvar. High-speed particle image velocimetry to assess cardiac fluid dynamics in vitro: From performance to validation. Eur. J. Mech. B/Fluids., 35:2–8, 2012.

16. O. Flores, L. Rossini, A Gonzalo, D. Vigneault, J. Bermejo, AM Kahn, E. McVeigh, M. Garcia-Villalba, and J C. del Alamo. Evaluation of blood stasis in the left atrium using patient-specific direct numerical simulations. In ERCOFTAC Workshop Direct and Large Eddy Simulation, pages 485–490. Springer, 2019.

17. Matthew D Ford, Gordan R Stuhne, Hristo N Nikolov, Damiaan F Habets, Stephen P Lownie, David W Holdsworth, and David A Steinman. Virtual angiography for visualization and validation of computational models of aneurysm hemodynamics. IEEE Trans. Med. Imaging., 24(12):1586–1592, 2005.

18. M. Garcia-Villalba, L. Rossini, A. Gonzalo, D. Vigneault, P. Martinez-Legazpi, E. Durán, O. Flores, J. Bermejo, E. McVeigh, and J. C. Kahn, A. M. del Alamo. Demonstration of patient-specific simulations to assess left atrial appendage thrombogenesis risk. Frontiers Physiol., 12:596596, 2021.

19. M. E. Goldman, L. A. Pearce, R. G. Hart, M. Zabalgoitia, R. W. Asinger, R. Safford, J. L. Halperin, Stroke Prevention in Atrial Fibrillation Investigators, et al. Pathophysiologic correlates of thromboembolism in nonvalvular atrial fibrillation: I. reduced flow velocity in the left atrial appendage (the stroke prevention in atrial fibrillation [spaf-iii] study). J. Am. Soc. Echocardiog., 12(12):1080–1087, 1999.

20. A. Gonzalo, M. García-Villalba, L. Rossini, E. Durán, D. Vigneault, P. Martínez-Legazpi, O. Flores, J. Bermejo, E. McVeigh, A. M. Kahn, and J. C. delálamo. Non-newtonian blood rheology impacts left atrial stasis in patient-specific simulations. Int. J. Numer. Method Biomed. Eng., 38(6):e3597, 2022.

21. Noelia Grande Gutiérrez, Mark Alber, Andrew M Kahn, Jane C Burns, Mathew Mathew, Brian W McCrindle, and Alison L Marsden. Computational modeling of blood component transport related to coronary artery thrombosis in kawasaki disease. PLoS Comput. Biol., 17(9):e1009331, 2021.

22. David Green. Coagulation cascade. Hemodial Int., 10(S2):S2–S4, 2006.

23. S. Hendabadi, J. Bermejo, Y. Benito, R. Yotti, F. Fernández-Avilés, J. C. delálamo, and S. C. Shadden. Topology of blood transport in the human left ventricle by novel processing of doppler echocardiography. Ann. Biomed. Eng., 41:2603–2616, 2013.

24. C. Hirsch. Numerical computation of internal and external flows. (2nd edition). Elsevier, 2007.

25. Matthew F Hockin, Kenneth C Jones, Stephen J Everse, and Kenneth G Mann. A model for the stoichiometric regulation of blood coagulation. J. Biol. Chem., 277(21):18322–18333, 2002.

26. K. Itô, P. Henry Jr, et al. Diffusion processes and their sample paths: Reprint of the 1974 edition. Springer Science & Business Media, 1996.

27. Guang-Shan Jiang and Chi-Wang Shu. Efficient implementation of weighted ENO schemes. J. Comput. Phy., 126(1):202–228, 1996.

28. K. C. Jones and K. G. Mann. A model for the tissue factor pathway to thrombin. II. A mathematical simulation. J. Biol. Chem., 269(37):23367–23373, 1994.

29. MR Kaazempur-Mofrad and CR Ethier. Mass transport in an anatomically realistic human right coronary artery. Ann. Biomed. Eng., 29:121–127, 2001.

30. JP Keener and James Sneyd. Mathematical physiology 1: Cellular physiology. Springer New York, NY, USA, 2009.

31. L. J. Kelsey, J. T. Powell, P. E. Norman, K. Miller, and B. J. Doyle. A comparison of hemodynamic metrics and intraluminal thrombus burden in a common iliac artery aneurysm. Int. J. Numer. Method Biomed. Eng., 33(5):e2821, 2017.

32. D. N. Ku. Blood flow in arteries. Ann. Rev. Fluid Mech., 29(1):399–434, 1997.

33. Andrew L Kuharsky and Aaron L Fogelson. Surface-mediated control of blood coagulation: the role of binding site densities and platelet deposition. Biophys. J., 80(3):1050–1074, 2001.

34. Jeffrey H Lawson, Michael Kalafatis, Shari Stram, and Kenneth G Mann. A model for the tissue factor pathway to thrombin. i. an empirical study. J Biol Chem, 269(37):23357–23366, 1994.

35. K. Leiderman and A. Fogelson. Grow with the flow: a spatial–temporal model of platelet deposition and blood coagulation under flow. Math Med Biol., 28(1):47–84, 2011.

36. K. Leiderman and A. Fogelson. An overview of mathematical modeling of thrombus formation under flow. Thromb. Res, 133:S12–S14, 2014.

37. Y. Li, O. Amili, S. Moen, P. Van de Moortele, A. Grande, B. Jagadeesan, and F. Coletti. Flow residence time in intracranial aneurysms evaluated by in vitro 4d flow mri. J. Biomech., 141:111211, 2022.

38. A. I. Lobanov and T. K. Starozhilova. The effect of convective flows on blood coagulation processes. Pathophysiol. Haemos. Thromb., 34(2-3):121–134, 2005.

39. RG Macfarlane. An enzyme cascade in the blood clotting mechanism, and its function as a biochemical amplifier. Nature, 202(4931):498–499, 1964.

40. G. Mareels, R. Kaminsky, S. Eloot, and P. R. Verdonck. Particle image velocimetry–validated, computational fluid dynamics–based design to reduce shear stress and residence time in central venous hemodialysis catheters. Asaio J., 53(4):438–446, 2007.

41. M. Moriche, O. Flores, and M. Garcia-Villalba. On the aerodynamic forces on heaving and pitching airfoils at low reynolds number. J. Fluid Mech., 828:395–423, 2017.

42. M. N. Ngoepe, A. F. Frangi, J. V. Byrne, and Y. Ventikos. Thrombosis in cerebral aneurysms and the computational modeling thereof: a review. Front. Physiol., 9:306, 2018.

43. M. J. O’Rourke, J. P. McCullough, and S. Kelly. An investigation of the relationship between hemodynamics and thrombus deposition within patient-specific models of abdominal aortic aneurysm. Proc. Inst. Mech. Eng. H., 226(7):548–564, 2012.

44. Mikhail A Panteleev, Mikhail V Ovanesov, Dmitrii A Kireev, Aleksei M Shibeko, Elena I Sinauridze, Natalya M Ananyeva, Andrey A Butylin, Evgueni L Saenko, and Fazoil I Ataullakhanov. Spatial propagation and localization of blood coagulation are regulated by intrinsic and protein c pathways, respectively. Biophys. J., 90(5):1489–1500, 2006.

45. Ahmed Qureshi, Maximilian Balmus, Dmitry Nechipurenko, Fazoil Ataullakhanov, Steven Williams, Gregory Lip, David Nordsletten, Oleg Aslanidi, and Adelaide De Vecchi. Left atrial appendage morphology impacts thrombus formation risks in multi-physics atrial models. In 2021 Computing in Cardiology (CinC), volume 48, pages 1–4. IEEE, 2021.

46. Gary E Raskob, Pantep Angchaisuksiri, Alicia N Blanco, H Buller, Alexander Gallus, Beverley J Hunt, Elaine M Hylek, Ajay Kakkar, Stavros V Konstantinides, Micah McCumber, et al. Thrombosis: a major contributor to global disease burden. Arterioscler Thromb Vasc Biol, 34(11):2363–2371, 2014.

47. N. Ratto, A. Tokarev, P. Chelle, B. Tardy-Poncet, and V. Volpert. Clustering of thrombin generation test data using a reduced mathematical model of blood coagulation. Acta Biotheor., 68(1):21–43, 2020.

48. V.L. Rayz, L. Boussel, L. Ge, J.R. Leach, A.J. Martin, M.T. Lawton, C. McCulloch, and D. Saloner. Flow residence time and regions of intraluminal thrombus deposition in intracranial aneurysms. Ann. Biomed. Eng., 38(10):3058–3069, 2010.

49. P. J. Roache. Perspective: a method for uniform reporting of grid refinement studies. J. Fluids. Eng., 116:405–413, 1994.

50. Stanley L Robbins and Ramzi S Cotran. Pathologic basis of disease. Saunders, 1979.

51. L. Rossini, P. Martinez-Legazpi, V. Vu, L. Fernandez-Friera, C. Perez del Villar, S. Rodriguez-Lopez, Y. Benito, M.-G. Borja, D. Pastor-Escuredo, R. Yotti, et al. A clinical method for mapping and quantifying blood stasis in the left ventricle. J. Biomech., 49(11):2152–2161, 2016.

52. Matthew K Runyon, Bethany L Johnson-Kerner, and Rustem F Ismagilov. Minimal functional model of hemostasis in a biomimetic microfluidic system. Angew. Chem., 116(12):1557–1562, 2004.

53. J. H. Seo, T. Abd, R. T. George, and R. Mittal. A coupled chemo-fluidic computational model for thrombogenesis in infarcted left ventricles. Amer. J. Physiol.-Heart Circul. Physiol., 310(11):H1567–H1582, 2016.

54. A. Steingoetter, D. Weishaupt, P. Kunz, K. Mäder, H. Lengsfeld, M. Thumshirn, P. Boesiger, M. Fried, and W. Schwizer. Magnetic resonance imaging for the in vivo evaluation of gastric-retentive tablets. Pharm. Res., 20:2001–2007, 2003.

55. M. Tomaszewski, K. Sybilski, P. Baranowski, and J. Ma lachowski. Experimental and numerical flow analysis through arteries with stent using particle image velocimetry and computational fluid dynamics method. Biocybern. Biomed. Eng., 40(2):740–751, 2020.

56. M. Uhlmann. An immersed boundary method with direct forcing for the simulation of particulate flows. J. Comput. Phys., 209(2):448–476, 2005.

57. R. Vanninen, H. Manninen, and A. Ronkainen. Broad-based intracranial aneurysms: thrombosis induced by stent placement. Am. J. Neuroradiol., 24(2):263–266, 2003.

58. C. Vlachopoulos, M. O’Rourke, and W. W. Nichols. McDonald’s blood flow in arteries: theoretical, experimental and clinical principles. CRC press, 2011.

59. Aaron M Wendelboe and Gary E Raskob. Global burden of thrombosis: epidemiologic aspects. Circulation research, 118(9):1340–1347, 2016.

60. David M Wootton and David N Ku. Fluid mechanics of vascular systems, diseases, and thrombosis. Annu. Rev. Biomed. Eng., 1(1):299–329, 1999.

61. Alireza Yazdani, He Li, Jay D Humphrey, and George Em Karniadakis. A general shear-dependent model for thrombus formation. PLoS Comput. Biol., 13(1):e1005291, 2017.

62. V. I. Zarnitsina, F. Ataullakhanov, A. I. Lobanov, and O. L. Morozova. Dynamics of spatially nonuniform patterning in the model of blood coagulation. Chaos, 11(1):57–70, 2001.

