## Appendices for "Efficient multi-fidelity computation of blood coagulation under flow"

### Supporting information

**S1 Appendix. Itô's Differentiation.** Consider the residence time $T = T(x(t), t)$ as a function of position, $x(t)$, and time, $t$. Assume that the position is a stochastic process that can be modeled as a Wiener process, such as $X_t \in N(0, t)$. Then, differentiating with Itô's lemma up to second order we obtain

$$dT = \frac{\partial T}{\partial t}dt + \frac{\partial T}{\partial x}dX_t + \frac{1}{2}\frac{\partial^2 T}{\partial t^2}dt^2 + \frac{1}{2}\frac{\partial^2 T}{\partial x^2}dX_t^2 + \frac{\partial^2 T}{\partial x \partial t}dtdX_t. \tag{22}$$

Since $T$ is the residence time, then $\frac{\partial T}{\partial t} = 1$, resulting in

$$dT = dt + \frac{\partial T}{\partial x}dX_t + \frac{1}{2}\frac{\partial^2 T}{\partial t^2}dt^2 + \frac{1}{2}\frac{\partial^2 T}{\partial x^2}dX_t^2 + \frac{\partial^2 T}{\partial x \partial t}dtdX_t. \tag{23}$$

Computing the average (i.e., the expected value, $E[-]$) and neglecting terms $O(dt^2)$ yields

$$dE[T] = dt + \frac{1}{2}\frac{\partial^2}{\partial x^2}E[T]dt, \tag{24}$$

where $E[dX_t] = dE[X_t] = 0$ and $E[dX_t^2] = dt$, (see [26]). Therefore, the total derivative of the mean residence time, $E[T] = \overline{t_R}$, can be expressed as

$$\frac{d\overline{t_R}}{dt} = 1 + \frac{1}{2}\frac{\partial^2}{\partial x^2}\overline{t_R}. \tag{25}$$

Proceeding in the same way but with the second order moment of the residence time $T^2$ (now considering $\partial T^2/\partial t = 2T \cdot \partial T/\partial t = 2T$), we obtain

$$\frac{d\overline{t_R^2}}{dt} = 2\overline{t_R} + \frac{1}{2}\frac{\partial^2}{\partial x^2}\overline{t_R^2}. \tag{26}$$

It is important to note that the PDE for $\overline{t_R}^2$ is not the same as the PDE for $\overline{t_R^2}$, as can be seen comparing

$$\frac{d}{dt}\overline{t_R}^2 = 2\overline{t_R}\frac{d\overline{t_R}}{dt} = 2\overline{t_R}\left(1 + \frac{1}{2}\frac{\partial^2}{\partial x^2}\overline{t_R}\right) \tag{27}$$

with eq. (26).

To calculate the spatio-temporal evolution of the variance $\sigma_T^2$, one may proceed as follows:

$$\sigma_T^2 = \overline{t_R^2} - \overline{t_R}^2 \quad \rightarrow \quad \frac{d\sigma_T^2}{dt} = \frac{d\overline{t_R^2}}{dt} - \frac{d\overline{t_R}^2}{dt}. \tag{28}$$

Substitution of equations (27) and (26) into (28) yields

$$\frac{d\sigma_T^2}{dt} = \frac{1}{2}\frac{\partial^2}{\partial x^2}\overline{t_R^2} - \frac{1}{2}\frac{\partial^2}{\partial x^2}\overline{t_R}^2 + \left(\frac{\partial \overline{t_R}}{\partial x}\right)^2 = \frac{1}{2}\frac{\partial^2}{\partial x^2}\sigma_T^2 + \left(\frac{\partial \overline{t_R}}{\partial x}\right)^2. \tag{29}$$

This expression shows that the variance grows with a source term proportional to the square of the spatial gradient of $\overline{t_R}$.

**S2 Appendix. Multi-Fidelity Model of Third Order.** Assuming that $g_i(T)$ is differentiable up to the third order, we can follow the same approach as described in section Higher-order Multi-Fidelity Approximations to derive the following expression:

$$u_i(\mathbf{x}, t) \approx g_i(\overline{t_R}) + g_i''(\overline{t_R})\frac{\sigma_T^2}{2} + g_i'''(\overline{t_R})\frac{\gamma_T}{3!} \tag{30}$$

where $\gamma = \int_{-\infty}^{\infty} (T - \overline{t_R})^3 f_T(T; \mathbf{x}, t) dT$. However, this third-order approximation introduces the third-order moment of the resident time at each fluid particle, which is denoted as $\overline{t_R^3}$. The spatio-temporal evolution of this term in an Eulerian framework can be obtained using a similar method as described in S1 Appendix for $\overline{t_R^2}$, yielding the following EDE:

$$\frac{D\overline{t_R^3}}{Dt} = 3\overline{t_R^2} + \frac{1}{2}\frac{\partial^2}{\partial x^2}\overline{t_R^3}. \tag{31}$$

Ignoring the diffusive term in (31) we obtain the *true* PDE for $\overline{t_R^3}$:

$$\frac{D\overline{t_R^3}}{Dt} = 3\overline{t_R^2}. \tag{32}$$

Consequently, the steps for the third order multi-fidelity approach are:

- **Third-order (MuFi-3)**:
  - Solve N ODEs eq. (6) to calculate $g_i(t)$ and its second and third temporal derivative $g_i''(t)$ and $g_i'''(t)$.
  - Solve three PDEs: Calculate $\overline{t_R}(\mathbf{x}, t)$ from eq. (4), $\overline{t_R^2}(\mathbf{x}, t)$ from (14) and $\overline{t_R^3}(\mathbf{x}, t)$ from (32).
  - Calculate $\sigma_T^2 = \overline{t_R^2} - \overline{t_R}^2$, $\gamma_T = \overline{t_R^3} - 3\overline{t_R}\sigma_T^2 - \overline{t_R}^3$, and map $u_i(\mathbf{x}, t) \approx g_i(\overline{t_R}) + g_i''(\overline{t_R})\frac{\sigma_T^2}{2} + g_i'''(\overline{t_R})\frac{\gamma_T}{3!}$.

**S3 Appendix. Womersley Inflow Boundary Condition.** Considering the unidirectional motion of an incompressible flow with density $\rho$ and kinematic viscosity $\nu$, between two parallel flat walls separated by a distance $H$ (as depicted in Figure 1), the velocity of the fluid ($v_x$) is described in time, by simplifying the Navier-Stokes equation as follows:

$$\frac{\partial v_x}{\partial t} = \nu \frac{\partial^2 v_x}{\partial y^2} - \frac{1}{\rho}\frac{\partial P}{\partial x}. \tag{33}$$

In this equation, $P$ is the pressure, while $x$ and $y$ are streamwise and wall-normal coordinates, respectively. Assume a pulsating pressure gradient,

$$\frac{\partial P}{\partial x} = \frac{\Delta P}{H} + \Re(A^* e^{i\omega t}), \tag{34}$$

where the operator $\Re()$ denotes the real part and $\Delta P$ is the average pressure gradient over one cycle. Then, a non-dimensional solution can be obtained for the flow with no-slip boundary conditions, i.e., $v_x(y = -H/2) = v_x(y = H/2) = 0$. This solution is characterized by velocity and length represented by $U_c$ and $H$, respectively, and can be determined by applying the equation (33).

$$\tilde{v}_x = \frac{Re}{2}\Delta\tilde{P}(1/4 - \tilde{y}^2) - \Re\left[i\frac{\tilde{A}^* Re}{\alpha^2}\left(\frac{e^{-\lambda H/2} - e^{\lambda H/2}}{e^{-\lambda H} - e^{\lambda H}}(e^{-\lambda H\tilde{y}} + e^{\lambda H\tilde{y}}) - 1\right)e^{i(\alpha^2/Re)\tilde{t}}\right], \tag{35}$$

where $\lambda = \sqrt{\frac{i\omega}{\nu}}$, $\Delta\tilde{P} = \rho U_c^2(\Delta P)$, $S_t = \omega H/U_c$, $\tilde{A}^* = A^* H/(\rho U_c^2 \Delta\tilde{P})$, $\tilde{t} = tU_c/H$ and Womersley number $\alpha = \sqrt{2\pi Re S_t}$, and Reynolds number $Re = U_c H/\nu$.

**S4 Appendix. Error maps for factor XIa and PCa.** For completeness, Figs. 13 and 14 presents the error maps for the concentrations of factor XIa and PCa, respectively. The errors in the concentration are plotted as a function of $\overline{t_R}$ and $\sigma_T^2$, analogous to Fig. 11 for the errors in the concentration of thrombine.

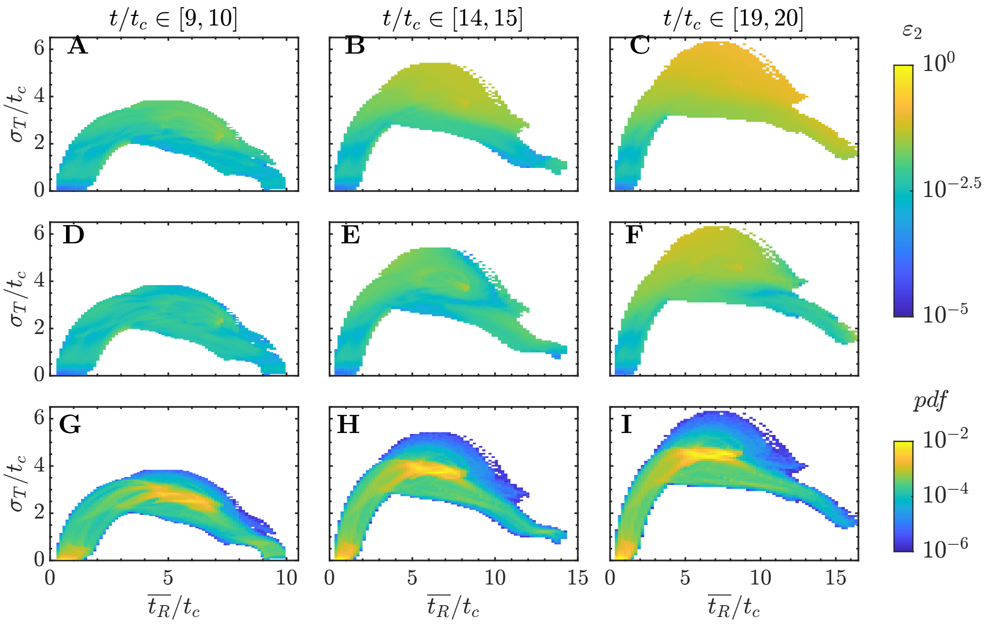


Figure 13: Error maps for factor XIa. A-C: Relative error in factor XIa concentration in MuFi-1, $\varepsilon_2^1$, as a function of residence time and its standard deviation. D-F: Same, but for MuFi-2, $\varepsilon_2^2$. G-I: Joint probability density function of residence time ($\overline{t_R}$) and its standard deviation ($\sigma_T$). Data for all panels is compiled inside cavity during three different cycles, as indicated on the top row. A,D,G: 10th cycle. B,E,H: 15th cycle. C,F,I: 20th cycle.

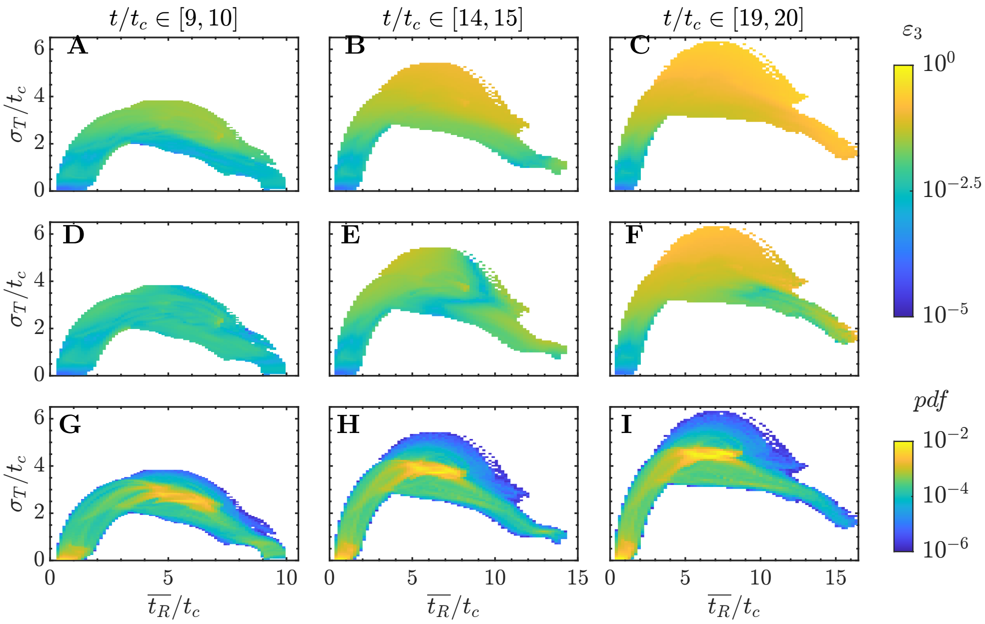


Figure 14: Error maps for protein C activated, PCa. A-C: Relative error in PCa concentration in MuFi-1, $\varepsilon_3^1$, as a function of residence time and its standard deviation. D-F: Same, but for MuFi-2, $\varepsilon_3^2$. G-I: Joint probability density function of residence time ($\overline{t_R}$) and its standard deviation ($\sigma_T$). Data for all panels is compiled inside cavity during three different cycles, as indicated on the top row. A,D,G: 10th cycle. B,E,H: 15th cycle. C,F,I: 20th cycle.
